# Reconciling time and prediction error theories of associative learning

**DOI:** 10.1101/2025.01.25.634891

**Authors:** Noé Hamou, Samuel J. Gershman, Gautam Reddy

## Abstract

Learning involves forming associations between sensory events that have a consistent temporal relationship. Influential theories based on prediction errors explain numerous behavioral and neurobiological observations but do not account for how animals measure the passage of time. Here, we propose a Bayesian theory for temporal causal learning, where the structure of inter-stimulus intervals is used to infer the singular cause of a rewarding stimulus. We show that a single assumption of timescale invariance, formulated as an hierarchical generative model, is sufficient to explain a puzzling set of learning phenomena, including the power-law dependence of acquisition on inter-trial intervals and timescale invariance in response profiles. A biologically plausible algorithm for inference recapitulates salient aspects of both timing and prediction error theories. The theory predicts neural signals with distinct dynamics that encode causal associations and temporal structure.

## I. INTRODUCTION

Animals learn the structure of a novel environment by forming associations between events that share consistent spatial and temporal relationships. Many principles of associative learning have been discovered within the classical conditioning paradigm, where learning is typically measured by an animal’s anticipatory response to a rewarding stimulus (US) that consistently follows a cue (CS) [1]. Classical conditioning experiments reveal a rich set of behavioral phenomena, including contingency degradation, blocking, and conditioned inhibition, which can be explained by reward prediction error (RPE) models. Notable examples include the Rescorla-Wagner model [2] and its temporal-difference (TD) generalizations [3, 4]. Neuroscientific studies provide strong support for RPE models, demonstrating that the dynamics of mesolimbic dopamine during learning and extinction match those of an RPE signal [5–11].

Classical RPE models do not easily explain how animals form associations across events separated by timescales spanning many orders of magnitude [12, 13]. TD models typically discretize time into states that tile the interval between the cue and reward. A TD learning rule sequentially propagates prediction errors backward in time along those discrete states, explaining how associations between distal cues and rewards could be learned [4, 14, 15]. The choice of discretization fixes an intrinsic timescale that governs the rate at which an association is acquired and the temporal precision available when anticipating reward.

However, a puzzling empirical observation is the absence of an intrinsic timescale that sets the rate of learning. Instead, the number of trials (*n*_acq_) required for an animal to exhibit an anticipatory response is primarily determined by the ratio of the cue-reward interval (*T*) to the reward-reward interval (*C*) (Figure 1a). Specifically, *n*_acq_ has an approximate power-law relationship with *C/T* [16, 17] (Figure 1c). Further, an animal’s anticipatory response exhibits Weber-law-like scaling [18], where response profiles across experiments collapse when time is rescaled by the cue-reward interval (Figure 1d). Other puzzling observations (from an RPE point of view) include discontinuous learning curves [19] (Figure 1b), the impact of reward pre-conditioning trials on acquisition (Figure 1e), and a variety of timing-related observations suggesting that animals explicitly encode intervals between events [20]. Recent neurobiological findings have further questioned whether mesolimbic dopamine encodes a pure RPE signal [21], reinvigorating efforts to develop a unified framework that reconciles RPE models with these phenomena.

**FIG. 1.**
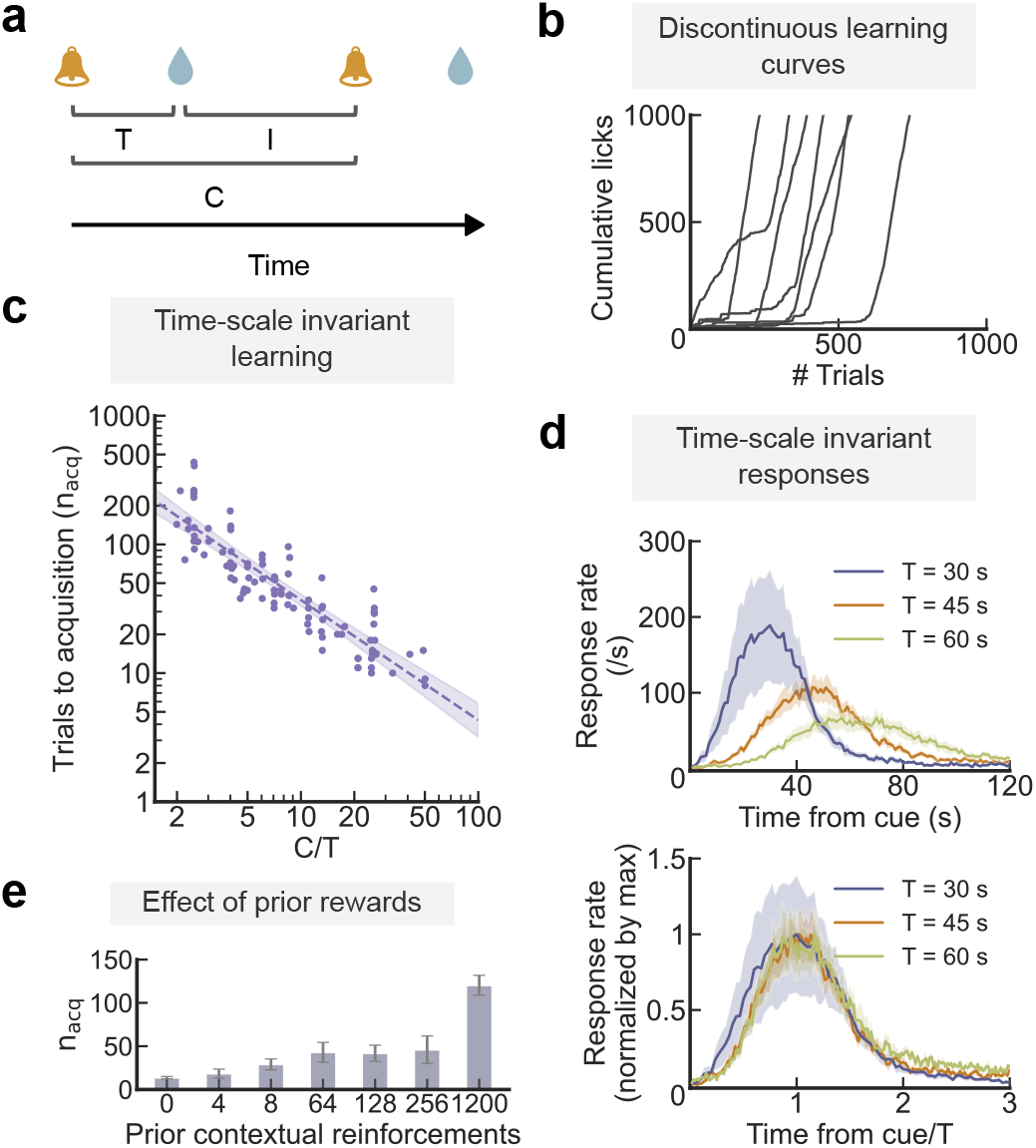
Learning phenomena in a classical conditioning paradigm. **(a)**: Schematic representation of the delay-conditioning protocol. A reward (such as water) is presented a fixed interval after the cue (such as a bell). The interval between the cue and the reward is *T*, between the reward and the next cue is *I*, while *C* = *I* + *T* is the the interval between consecutive cues/rewards. **(b)**: Discontinuous learning curves. Animals trained in a Pavlovian conditioning framework learn to anticipate rewards by licking in response to a cue. Cumulative lick counts are displayed, with traces shown up to the 1000th lick per animal. Data adapted from [21]. **(c)**: Timescale invariant learning. The number of trials required for acquisition *n*_acq_ is plotted against the *C/T* ratio on a logarithmic scale. Data are compiled from studies across different laboratories, as listed in [16, 17]. A linear fit (dashed line) is shown, highlighting the power-law relationship between *n*_acq_ and *C/T*. **(d)**: Timescale invariance in response profiles. In the top panel, response rates are plotted for various cue-reward intervals *T* (left). In the bottom panel, both response rates and time are normalized by their respective maxima for each *T*. Data are adapted from [18] and sourced from the *Timing Database* [32]. **(e)** Effect of prior rewards. During pre-conditioning with rewards, animals receive prior contextual reinforcements before any cues are introduced. Increasing the number of contextual trials results in animals requiring more cues to form the association. Data from [33, 34].

Alternative models have been proposed to account for some of these phenomena [14, 16, 21–27] (discussed further in Appendix C). Discontinuous learning curves can be explained based on the nature of TD signal propagation in structured environments [14], or by assuming that animals implement approximate Bayesian inference by implementing a sampling algorithm [28]. Building on rate estimation theory (RET) [16], a line of work [22, 29] argues that the rate of acquisition is determined by the additional information (in bits) the cue provides about reward timing relative to the background context, which depends on *C/T*. A recent model grounds RET in learning theoretic terms and makes a link with RPE-like models [25]. Another recent theoretical framework, called retrospective causal learning theory (RCT) [21], proposes that animals learn causal associations using an eligibility trace mechanism whose characteristic timescale is set by the inter-trial interval. RET and RCT help rationalize why *n*_acq_ depends primarily on the ratio *C/T*. Other models invoke cue competition [23] to highlight the influence of reward pre-conditioning, and predictive representations of stimulus-reward intervals [12, 26, 30, 31] to explain how an animal could form associations across multiple timescales.

Here, we show that prior models describing complementary aspects of associative learning can be synthesized within one common framework. The framework can be viewed as a version of model-based RL where learning temporal structure plays a central role. We present two main contributions. First, we formulate a general Bayesian framework for timing-based causal learning that describes how causal associations are learned and how these associations determine anticipatory responses. The inference process involves estimating two interdependent quantities: the distribution of intervals between stimuli, and a probabilistic measure of the causal association between them. The key insight is that a single assumption of timescale invariance, formulated as an hierarchical generative model, quantitatively explains the phenomena described in Figure 1. Second, we show that online algorithms for learning distributions of stimulus-reward intervals closely resemble prediction error models. We propose a novel learning rule for estimating causal associations between stimuli, which predicts neural signals with distinctive dynamics that encode causal associations. When applied to a common classical conditioning protocol (Figure 1a), the model reproduces the core features of the Rescorla-Wagner model, including contingency degradation, blocking, extinction and a prediction error signal consistent with an RPE.

## II. RESULTS

### A. A Bayesian framework for timing-based causal learning

We consider a scenario where the animal predicts *when* a rewarding stimulus (*r*) will appear based on the timing of past stimuli (Figure 2a). The rewarding stimulus *r* appears at some (possibly stochastic) interval after a stimulus that causes it. The causal stimulus *c* may either be a previous occurrence of *r* itself or a previous occurrence of a stimulus *c* amongst a set of possible non-rewarding stimuli (Figure 2a). Our theory has two main features: (1) that the animal estimates the likelihood that one of the stimuli causes *r* based on statistical regularities in stimulus-reward intervals, and (2) that the animal displays an anticipatory response that maximizes long-term reward. The anticipatory response relies on a predictive map of when *r* will occur next given the historical record of when past stimuli have occurred.

**FIG. 2.**
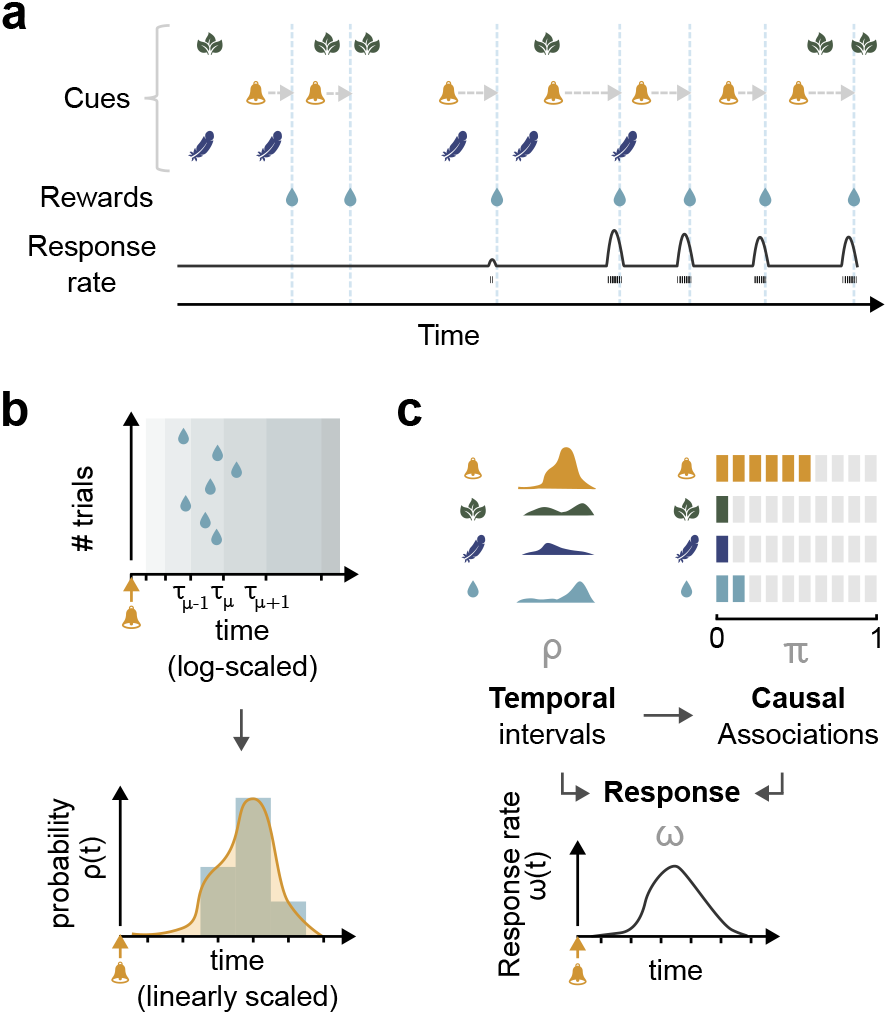
Overview of the model. **(a)** Schematic representation of the timing-based causal learning framework. Stimuli are point-events in time. Based on the timing between stimuli and rewards, the model agent learns to respond in anticipation of reward following stimuli that are predictive of rewards. In this example, the orange bell is the best predictor of when reward will occur. **(b)** (Top) Schematic representation of the learned histogram of the interval between the causal stimulus and reward. Rewards are placed in bins, where bin locations *τ*_1_, *τ*_2_, …, *τ*_*K*_ are uniformly spaced on a logarithmic time axis. (Bottom) The corresponding distribution of intervals (yellow) between the causal stimulus and the reward is obtained by smoothening the histogram (blue). **(c)** Algorithmic steps of the proposed Bayesian learning theory. The agent learns to estimate distributions of intervals between each stimulus and reward (including the distribution of intervals from reward to reward). The agent uses these estimates of interval distributions to estimate the probability that each of the stimuli causes reward. The agent then combines the estimated interval distribution and causal associations to produce an anticipatory response to the reward.

The framework is cast as an hierarchical Bayesian model. Given the historical record until time *t*, Bayesian inference lets us compute the probability density that *r* will appear at time *t*. This probability is given by

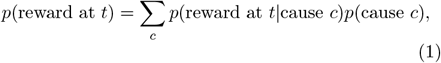

which we express as *p*(*t*) = ∑_*c*_ *ρ*_*c*_(*t*)*π*_*c*_(*t*), where the sum over *c* includes *r* and all possible non-rewarding stimuli in the context. *ρ*_*c*_(*t*) encodes the information the animal has acquired about the distribution of intervals between *c* and *r. π*_*c*_(*t*) is the *association*, defined as the posterior probability that *c* causes *r*. Using Bayes’ rule, the association *π*_*c*_(*t*) is proportional to the likelihood of observing the historical record until time *t* if *c* were the cause, weighted by the prior probability that *c* is the cause. The association can generally be expressed as

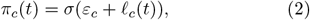

where *ε*_*c*_ is the relative log prior, 𝓁_*c*_(*t*) is the relative log likelihood and 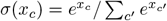 is the softmax function. *ε*_*c*_ and 𝓁_*c*_ are respectively measured relative to the log prior and log likelihood that reward causes reward 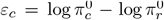 and 𝓁_*c*_(*t*) = log *ρ*_*c*_(*t*) − log *ρ*_*r*_(*t*)). The possibility that reward causes reward plays a similar role as a common assumption in prior models that a non-rewarding stimulus competes with a background contextual cue [35, 36]. The relative log prior *ε*_*c*_ captures a notion of “preparedness” [37], that is, the propensity for a particular stimulus to be associated with the reward based on the animal’s past experiences or innate biases. We now expand on the theory’s two key features, timescale invariance and reward maximization.

#### Timescale invariance

Timescale invariance is motivated by the viewpoint that an animal could form associations between contiguous events (separated by fractions of seconds) but also between distant events (separated by minutes or hours). One would expect a representation of time intervals that supports associations across timescales separated by orders of magnitude to be logarithmic. We will show that this assumption is also sufficient to explain experimental data. Specifically, we assume the interval distribution between the causal stimulus and reward is represented as a (smoothed) histogram, where the temporal locations of the *K* histogram bins are *τ*_1_, *τ*_2_, *τ*_3_, …, *τ*_*K*_ (Figure 2b). The animal learns this smoothed histogram during conditioning. Importantly, the timescales are spaced uniformly on a logarithmic scale, *τ*_*µ*+1_ − *τ*_*µ*_ = *kτ*_*µ*_ for *k* ≪ 1.

This formulation of timescale invariance can be expressed mathematically as a generative model using a Dirichlet-multinomial distribution compounded with a scale-invariant emission function (Methods). The complete inferential framework is expressive enough to allow for multi-modal distributions of stimulus-reward intervals and partial reinforcement. Exact inference can be computationally hard in certain scenarios due to the many possible assignments between rewards and their causes when stimuli and rewards are interleaved. We discuss approximate algorithms for inference in Appendix A, noting however that the assignment problem is absent for the delay-conditioning protocol considered here.

In this model, the probability density that the reward will occur at *t* if the causal stimulus *c* appeared at *t* – *δ* is given by

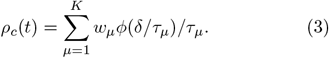

Intuitively, *w*_*µ*_ represents the probability that the reward will fall in the bin at timescale *τ*_*µ*_ and *ϕ* is a normalized basis function which smooths the estimated histogram (Figure 2b). The weights *w*_*µ*_ thus encode information about the distribution of stimulus-reward intervals. A uniform prior over *w*_*µ*_ leads to a 1*/δ* power-law prior distribution over stimulus-reward intervals. This scaling relation implies that shorter intervals are more likely, and that the relative likelihood of observing two intervals is equal to the ratio of those intervals.

#### Reward maximization

When an animal experiences a reward-predictive stimulus, it displays an anticipatory response (for example, by licking a water port) at a rate *ω*(*t*) that reflects its estimate of whether the reward will occur at time *t* [38, 39]. We show that if each response has a rate-dependent cost and future rewards are discounted at a discount rate *λ*, the optimal anticipatory response rate *ω*^*^(*t*) generally takes the form

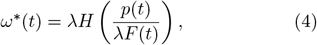

where 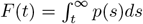 and *H* is a monotonic function (see Methods for the derivation). For example, if the cost of a response is independent of the rate *ω*, we find 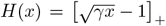 up to a maximal response rate. *γ* is a constant that depends on the subjective value the animal receives from the reward relative to the cost of each response.

Equation (4) implies that any threshold criterion applied on the response rate to deem that the animal has acquired the association is equivalent to a criterion on the certainty with which reward is predicted to occur, that is, *p*(*t*)*/F* (*t*) *>* Θ for some threshold Θ. Importantly, since *p*(*t*) depends on the product of *ρ*_*c*_(*t*) and *π*_*c*_(*t*), equation (4) further highlights that an association is acquired when the stimulus-reward interval is learned (*ρ*_*c*_ is sharply distributed) *and* when the stimulus is deemed causal (*π*_*c*_ ≈ 1) (Figure 2c).

### B. Timescale invariance explains timing-related phenomena

We now examine the behavior of the model when applied to the commonly used delay-conditioning protocol shown in Figure 1a. The experiment involves one unrewarding cue (CS) and a reward (US). The cue-reward and reward-reward intervals are fixed at *T* and *C*, respectively. The experiment begins with a pre-conditioning phase where the reward is delivered alone. The cue is introduced after *n*_*p*_ prior presentations of the reward. As in experiments, an association is deemed to be acquired when the rate of anticipatory response crosses a threshold.

Simulations successfully recapitulate discontinuous learning curves (Figure 3a) and timescale invariance in response profiles (Figure 3b). That is, response profiles across simulations with different *T* collapse when time since cue presentation is re-scaled by *T* and their amplitude is re-scaled by the maximum response value. Next, we examine how the number of trials for acquisition, *n*_acq_, depends on *n*_*p*_, *C* and *T*. We find that *n*_acq_ depends only on the ratio *C/T*. In particular, *n*_acq_ has an approximate power-law dependence on *C/T*, but tapers off for large values of *C/T* (Figure 3c). The relative log prior *ε*_*c*_ has a weak influence on *n*_acq_ and *C/T*. As noted previously [23, 33, 34], the pre-conditioning phase has a strong influence on *n*_acq_ in experiments (Figure 1e). This dependence is also captured by our model (Figure 3d, S1a).

**FIG. 3.**
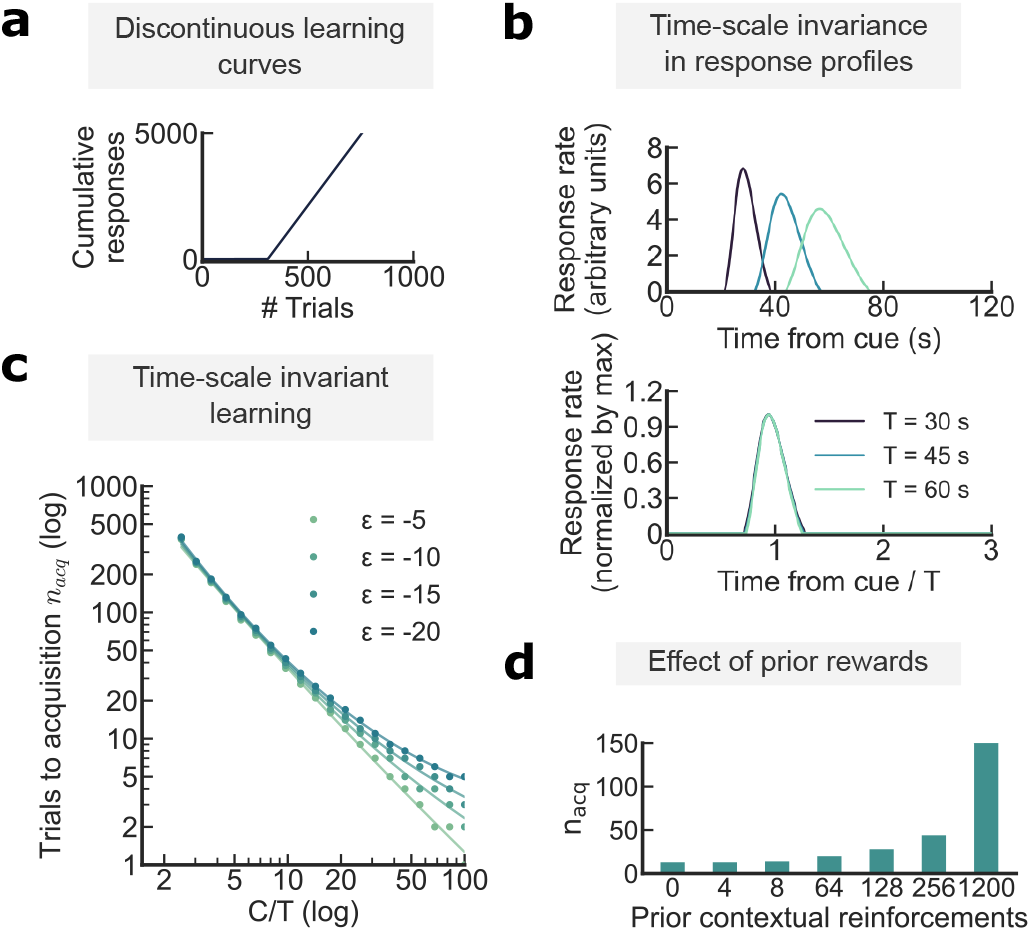
The Bayesian model reproduces timing-related phenomena. **(a)** Discontinuous learning curves. In accordance with Figure 1b, we plot the cumulative responses of a Bayesian agent during the delay-conditioning protocol described in Figure 1a. **(b)** Timescale invariance of response profiles. Response rates of the Bayesian agent for different values of cue-reward interval *T* (top) and after normalizing the response rate by the maximum response rate and re-scaling time by *T* (bottom).**(c)** Timescale invariant learning. The number of trials to acquisition, *n*_acq_, for the Bayesian agent are plotted against *C/T* for different values of the relative log-prior (*ε*_*c*_). Note the log-log scale. **(d)** Effect of prior rewards. In the Bayesian model, increasing the number of contextual trials results in the agent requiring more cues to form the association. See also Figure S1a.

A mathematical analysis of the model shows that timescale invariance in response profiles (Figure 3b), the power-law scaling of *n*_acq_ with respect to *C/T* (Figure 3c) and discontinuous learning curves (Figure 3a) are generic consequences of timescale invariance of the stimulus-reward interval distribution. We summarize the main results derived from our analysis and refer to the Methods for mathematical details.

Acquisition can only occur once the relative log likelihood 𝓁_*c*_ that the cue causes reward exceeds the relative log prior *ε*_*c*_, 𝓁_*c*_ *>* −*ε*_*c*_ (equation (2)). We show that 𝓁_*c*_ has a non-monotonic dependence on the number of presented cues: starting from zero, it first declines and subsequently rises to a positive value (Figure S1). The initial drop in 𝓁_*c*_ is due to the agent’s greater confidence in the reward-reward interval distribution acquired during the pre-conditioning phase. Longer pre-conditioning leads to a larger initial drop, which in turn leads to the significant dependence of *n*_acq_ on the number of prior rewards *n*_*p*_.

Since shorter intervals are more likely, the shorter cue-reward interval (*T < C*) leads to a subsequent rapid rise in 𝓁_*c*_ after a sufficient number of cue presentations. This rapid rise together with the sigmoidal dependence of the association *π*_*c*_ on 𝓁_*c*_ (equation (2)) leads to an abrupt learning of the cue-reward association. If the threshold criterion on the response rate for acquisition is small, then acquisition immediately follows. Thus, the theory suggests that acquisition is primarily limited by the time taken for the animal to establish that the cue is causal rather than the time taken for the animal to fully learn the cue-reward interval distribution (with some dependence on the acquisition criterion).

After the cue-reward association and interval are learned, the learned cue-reward interval distribution converges to *ρ*_*c*_(*δ*) ≈ *ϕ*(*δ/T*)*/T* (equation (3)). Since the response rate depends monotonically on *ρ*_*c*_, re-scaling *δ* with *T* and re-scaling the amplitude of the profile with the maximal value leads to timescale invariance in the response profile. The shape of this invariant response profile is determined primarily by the basis function *ϕ* and the response function *H*. Timescale invariance in the response profile is not exact in our model unless *H* is linear; however, the approximation is excellent despite the nonlinear *H* used in simulations (Figure 3c).

The analytical dependence of *n*_acq_ on *C* and *T* is in general non-trivial to obtain. We derive expressions for different parameter ranges (Methods). When *n*_*p*_, *K* ≫ *n*_acq_ ≫ 1 in particular, we find

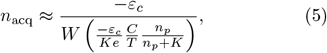

where *W* is the Lambert W function. *W* (*x*) ≈ *x* for *x* ≪ 1 implies *n*_acq_ ∝ (*C/T*)^−1^ whenever the argument in the *W* function of equation (5) is small.

To explain this reciprocal dependence of *n*_acq_ on *C/T*, we first note that *n*_acq_ depends on the log likelihood ratio log(*ρ*_*c*_(*T*)*/ρ*_*r*_(*C*)). Timescale invariance implies *ρ*_*c*_(*T*) ∝ 1*/T* and *ρ*_*r*_(*C*) ∝ 1*/C*, and thus *n*_acq_ depends only on the ratio *C/T*. By itself, however, this argument would imply a linear scaling of the evidence (∼*n* log *C/T*) after *n* cue-reward presentations. This linear scaling leads to an *n*_acq_ ∝ (log *C/T*)^−1^ relation inconsistent with data. The *n*_acq_ ∝ (*C/T*)^−1^ relation comes about because the animal’s estimate of the cue-reward interval also gets sharper with *n*. Specifically, *ρ*_*c*_(*T*) at the beginning of learning increases linearly with *n*: *ρ*_*c*_(*T*) ∝ *n/T*. The reward-reward interval is learned during the pre-conditioning phase, so that *ρ*_*r*_(*C*) = 1*/C*. Acquisition follows soon after the likelihood that the cue is causal exceeds the likelihood that the reward is causal *ρ*_*c*_(*T*) ≈ *ρ*_*r*_(*C*), which leads to *n*_acq_ ∝ (*C/T*)^−1^.

Our assumption that animals learn the *empirical histogram* of stimulus-reward intervals is important to explain the relationship between *n*_acq_ and *C/T*. To emphasize this point, we repeat the above analysis supposing the animal learns the rate parameter (drawn from a scale invariant prior) of a Gamma distribution. We find that while *n*_acq_ depends only on the ratio *C/T*, the specific relation is inverse logarithmic *n*_acq_ ∝ (log *C/T*)^−1^ (Methods).

### C. A theory of temporal causal learning

Building on the Bayesian theory, we now derive a biologically plausible algorithm for inference, which we call temporal causal learning (TCL). TCL involves two mechanisms: an update rule for learning stimulus-reward interval distributions, and an update rule for learning causal associations.

Learning the interval distribution involves updating the stimulus-specific weights *w*_*µ*_ corresponding to each timescale *τ*_*µ*_ (equation (3)). For each stimulus, we consider the update rule

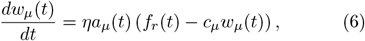

where *η* is a learning rate and *c*_*µ*_ is a constant that ensures the weights are normalized (Methods). *f*_*r*_(*t*) = ∑_*i*_ *δ*(*t* − *t*_*i*_) represents the reward signal, where the *t*_*i*_’s correspond to the times when the reward appeared in the past. *a*_*µ*_(*t*) is a stimulus-specific gating signal that determines which *w*_*µ*_ is updated when the reward appears (Figure S2b). The predicted probability that the reward will appear at time *t* is given by a linear readout 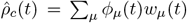, where *ϕ*_*µ*_ are normalized basis functions. The *a*_*µ*_s are set to zero after the reward appears until the next appearance of the stimulus.

With appropriate constraints on *c*_*µ*_, the gating signal *a*_*µ*_ and the basis functions *ϕ*_*µ*_, we show that equation (6) is a generic online kernel density estimation algorithm for learning distributions of intervals between two events (Methods). The kernel is specified by the choice of *a*_*µ*_ and *ϕ*_*µ*_. The update rule is consistent with an interpretation of the weight *w*_*µ*_ as encoding the estimated probability that the reward appears within the interval (*τ*_*µ*_, *τ*_*µ*+1_). The sum ∑_*µ*_ *w*_*µ*_ in turn encodes the probability that reward does indeed appear after the stimulus (∑_*µ*_ *w*_*µ*_ *<* 1 in a partial reinforcement paradigm).

We now show how the general update rule (6) can be used to derive a timescale invariant density estimator implementable in biological networks. Specifically, the gating signals *a*_*µ*_s are derived from a set of eligibility traces *ψ*_*µ*_s associated with each stimulus (Figure S2a). The eligibility traces for each stimulus (say *c*) are updated as

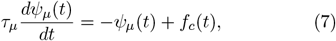

where *f*_*c*_(*t*) represents the stimulus train corresponding to stimulus *c*. A downstream network implements a soft winner-take-all operation, 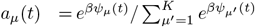. In the *β* → ∞ limit, we find that Δ*τ*_*µ*_ = *kτ*_*µ*_ and *k* ≪ 1 imply *a*_*µ*_(*t*) ≈ 1 in the interval (*τ*_*µ*_, *τ*_*µ*+1_) after the stimulus and 0 otherwise. Thus, *a*_*µ*_(*t*) represents the activity of “time cells” that are active in the interval *τ*_*µ*_ to *τ*_*µ*+1_ after the stimulus is presented (Figure S2b). In simulations, we use normalized gamma functions with scale parameter *τ*_*µ*_ and a fixed shape parameter as basis functions *ϕ*_*µ*_.

To derive an update rule for learning causal associations, we observe that the log likelihood *L*_*n*_ after *n* trials can generally be written as *L*_*n*_ = log *P* (data at *n* past data) + *L*_*n*−1_. Based on this recursive equation, we propose an update rule for the relative log likelihood (𝓁_*c*_) that the stimulus *c* is causal:

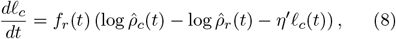

where *η*^*′*^ is a small constant that determines how many past events are averaged over when estimating 𝓁_*c*_. The pre-factor *f*_*r*_(*t*) in equation (8) indicates that the causal association is updated whenever the reward appears.

### D. TCL reproduces both prediction-error and timing-related phenomena

The model when applied to the delay-conditioning protocol reproduces discontinuous learning curves and timescale invariance in response profiles (Figure S3). The model also recapitulates the approximate power-law scaling of *n*_acq_ with *C/T* and has an excellent match with the data (Figure 4). All simulations of the model used the same set of parameters (see Methods). Notably, the approximate power-law behavior is preserved across a broad range of model parameters (Figure S4).

**FIG. 4.**
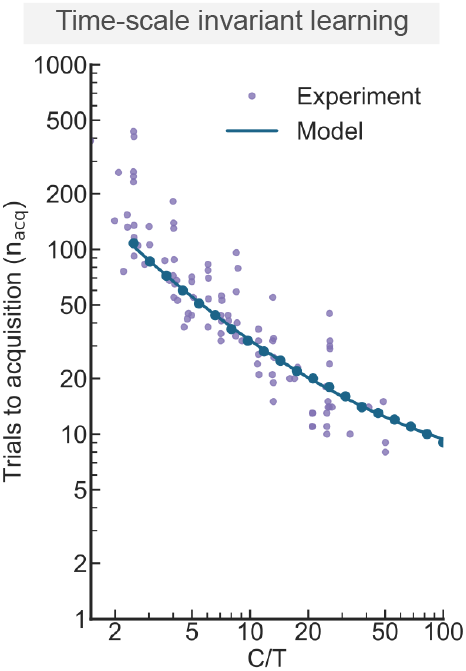
The online approximation version of our model (TCL) reproduces timescale invariant learning. Timescale invariant learning in the online model and in experiments [17]. The experimental points (purple dots) correspond to the data shown in Figure 1c. The model was trained on the delay-conditioning protocol described in Figure 1a with varying *C/T* ratios.

Observing that the term in the parenthesis in equation (6) resembles a prediction error, we hypothesized that TCL can reproduce phenomena attributed to RPE learning, such as extinction (Figure 5a,b), blocking (Figure 5c) and contingency degradation (Figure 5d). Indeed, in our model, we observe extinction of an acquired response when a previously expected reward is omitted (Figure 5a). Consistent with experiments [40], the rate of extinction is independent of the *C/T* ratio (Figure 5b). The TCL model also successfully captures the blocking effect (Figure 5c). Specifically, simulations show that the acquisition of a new stimulus-reward association is impaired when the reward has already been paired with a different stimulus, aligning with the classical blocking phenomenon. Finally, the TCL model replicates the contingency degradation effect. Contingency degradation is the reduction of an animal’s anticipatory response when additional uncued rewards are introduced after the cue-reward association is learned [41–43]. This effect arises in our model because the introduction of new rewards in the inter-trial period shortens the intervals between rewards, thus increasing the likelihood that rewards are caused by past rewards rather than past cues (Figure 5d).

**FIG. 5.**
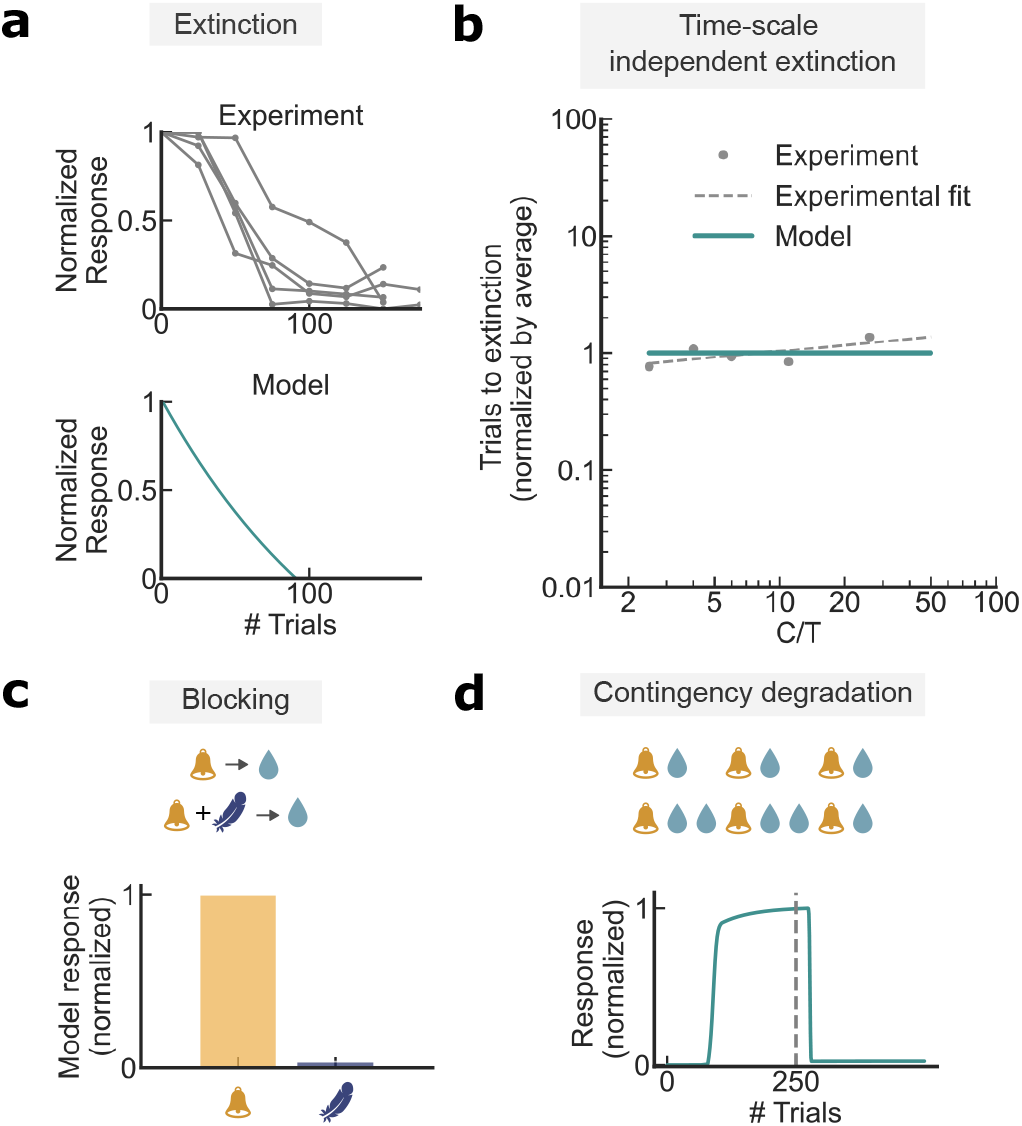
The TCL model is consistent with classical conditioning results. **(a)** Extinction in experiments (top) and TCL simulations (bottom). Individual curves correspond to individual mice. The normalization is by the maximum response rate. Data obtained from [21]. **(b)** Extinction rates are not dependent on *C/T* in experiments and TCL simulations. Data obtained from [40]. **(c)** Schematic representation of the blocking paradigm and the blocking effect in the TCL model. The response was normalized by the maximum response out of the two stimuli. **(d)** Schematic representation of the contingency degradation paradigm and of contingency degradation in the online model. The dashed gray line indicates when contingency degradation starts, which corresponds to the trial at which additional rewards are introduced in-between cue presentations. The response is normalized by the maximum response.

### E. The dynamics of neural correlates during learning

The dynamics of the quantities related to learning intervals (Δ*w*_*µ*_, *w*_*µ*_), associations (Δ𝓁_*c*_, 𝓁_*c*_, *π*_*c*_) and response (*ω*) are shown in Figure 6 for the delay-conditioning protocol. The update rule for Δ*w*_*µ*_ displays similar behavior as an RPE signal on the delay-conditioning protocol, though their behavior may differ when applied to other protocols. Specifically, the appearance of a reward at a certain interval triggers the positive update of weights corresponding to that interval, which eventually decays to zero while the interval distribution is learned (Figure 6a). The absence of rewards after the cue-reward interval has been learned leads to a concomitant negative update.

**FIG. 6.**
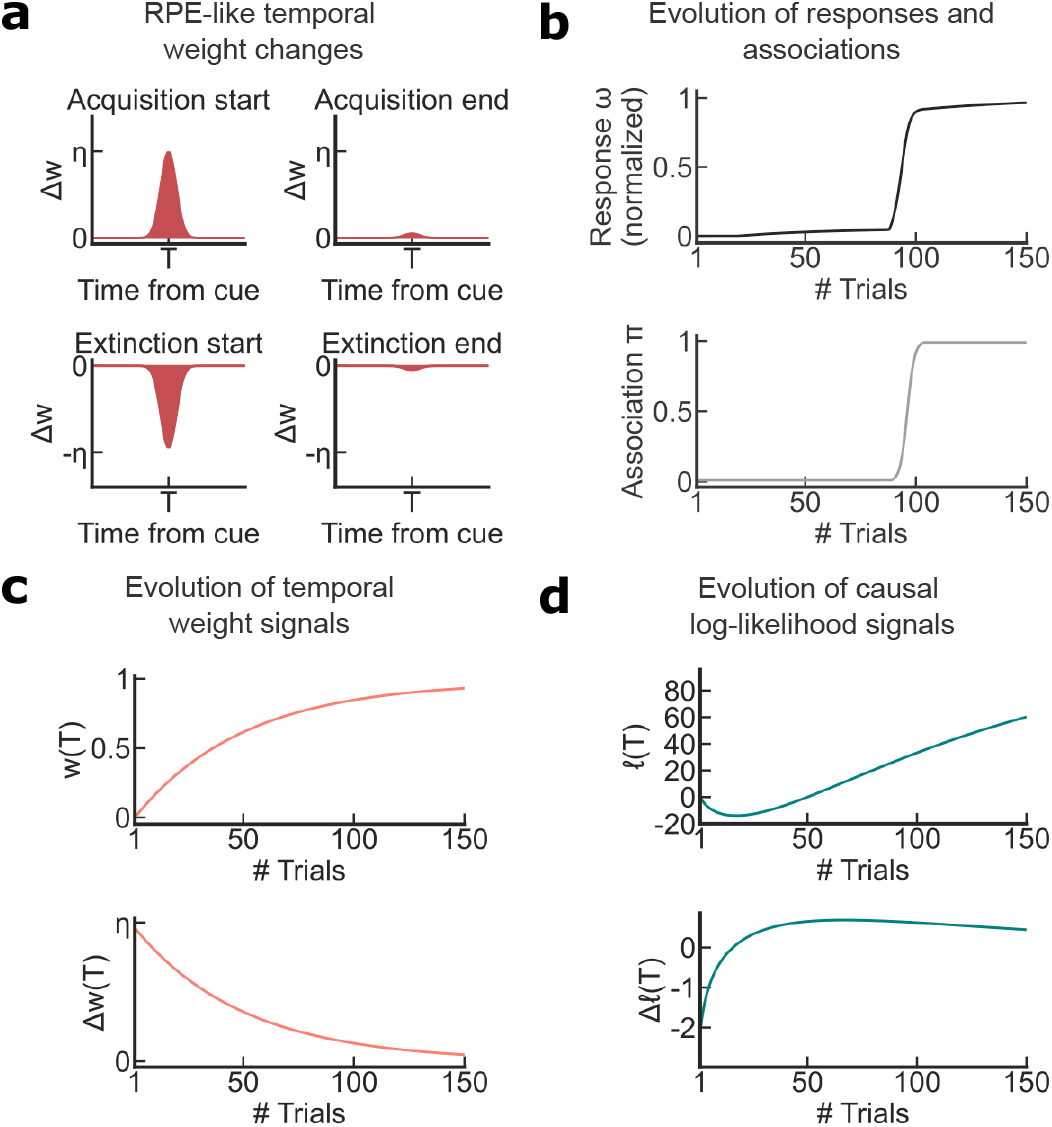
The TCL model predicts the existence of two learning signals: a classical RPE-like signal and a novel causal association signal. **(a)** RPE-like change of temporal weights during acquisition (top) and during extinction (bottom) **(b)** Evolution of the response (top) and of the association (bottom) across learning. **(c)** Evolution of the weights (top) and of the change in weights (bottom) across learning. **(d)** Evolution of the relative log-likelihood (top) and of the change in relative log-likelihood (bottom) across learning.

In the Rescorla-Wagner model, the response is considered to be a direct reflection of the associative strength between the cue and reward. Our model aligns with this picture; the abrupt acquisition of the response coincides with the acquisition of the association (Figure 6b). However, since the association increases together with the weights, the acquisition of the response will also be correlated with the weights that encode timing information (Figure 6c). Thus, whether a neural signal encodes causal associations or timing could be challenging to disentangle in experiments.

The update rule for the relative log likelihood (equation 8) predicts a non-monotonic reward-triggered signal, with the magnitude of the update peaking just before acquisition (Figure 6d). The magnitude of the negative dip in 𝓁_*c*_ increases with the number of prior rewards presented during the pre-conditioning phase. The response is acquired soon after 𝓁_*c*_ becomes positive. We note however that equation 8 is not the unique update rule that recapitulates this phenomenology. For example, it is possible that the log likelihoods for each stimulus (rather than *relative* log likelihood) are represented independently, which are later mixed when determining the response.

## III. DISCUSSION

A longstanding puzzle is the absence of a fixed intrinsic timescale for how quickly animals acquire associations. Curiously, the inter-trial interval has a large influence on learning rate: scaling the inter-trial interval by a factor of ten reduces the number of trials required for acquisition (*n*_acq_) by approximately the same factor [16, 29, 44]. We propose a Bayesian causal learning framework to address this puzzle and other unresolved learning phenomena that are not easily explained by existing models. Our approach synthesizes features of prior models into one framework, and highlights the central role played by temporal structure learning for forming associations.

The key insight is that a single assumption of timescale invariance, formulated in terms of how animals represent and learn distributions of stimulus-reward intervals when maximizing reward, can quantitatively account for abrupt learning curves, timescale invariance in response profiles and the quantitative relationship between *n*_acq_ and the ratio of the reward-reward and cue-reward intervals (*C/T*).

Guided by the intuitive notion that animals form associations across many timescales, we propose that animals learn kernel density estimates of interval distributions while measuring time on a logarithmic scale. If all intervals on this scale are equally likely, then the probability of observing a stimulus-reward interval is inversely proportional to that interval. Using a reward maximization framework to link anticipatory response with the animal’s temporal predictive map, we show that timescale invariance in interval distributions naturally leads to a Weber law scaling in response profiles. Further, since the likelihood of observing an interval is inversely proportional to the interval, the evidence (i.e., relative log likelihood) that the cue causes reward, and thus the number of trials to acquisition (*n*_acq_), depends only on *C/T*.

The theory predicts a specific non-trivial relationship between *n*_acq_ and *C/T* that depends both on the animal’s prior probability of forming the cue-reward association and on the animal’s exposure to the reward prior to cue-reward pairing. For a broad parameter range, we show that this relation approximates the empirically observed power-law relation between *n*_acq_ and *C/T*, but we expect deviations from this law, particularly when *C/T* is large. The non-trivial relation between *n*_acq_ and *C/T* arises because the evidence that the cue is causal increases supralinearly (∼*n* log(*nC/T*)) with the number of cue-reward presentations (*n*). This supralinear relation combined with the nonlinear relationship between evidence and response produces an abrupt learning effect akin to an “a-ha” moment when the evidence overcomes the prior. We predict that the relationship between *n*_acq_ and *C/T* switches from the approximate *n*_acq_ ∝ (*C/T*)^−1^ scaling observed in experiments to an *n*_acq_ ∝ (log *C/T*)^−1^ dependence (and a larger learning rate overall) if the animal is not significantly pre-conditioned to rewards.

Building on the Bayesian theory, we propose a biologically plausible model of inference, which we call temporal causal learning (TCL, Figure 7). The TCL update rule for learning intervals (equations (6), (7)) is closely related to a line of work highlighting the role of time cells and temporal context cells in associative learning [24, 45–47]. As in these models, *a*_*µ*_ reflects the activity of time cells and *ψ*_*µ*_ (the eligibility trace) reflects the activity of temporal context cells. However, the TCL update rule has certain key differences. First, we show that time cells can be derived from eligibility traces using a simple winner-take-all circuit rather than an approximate Laplace transform [24]. Next, our update rule has a precise interpretation as an online kernel density estimator for learning interval distributions, where the kernel is specified by *a*_*µ*_ and *ϕ*_*µ*_. This connection to density estimation emphasizes that there are multiple update rules, corresponding to different choices of the kernel, that can approximate inference and explain data equally well. Thus, our proposed algorithm using eligibility traces and a downstream winner-take-all circuit is one of potentially many biologically plausible mechanisms for implementing temporal causal learning. Finally, the correspondence with the Bayesian framework highlights the need for another update rule (equation 8) to learn causal associations and a reward maximization principle to connect interval estimation with response (equation 4).

**FIG. 7.**
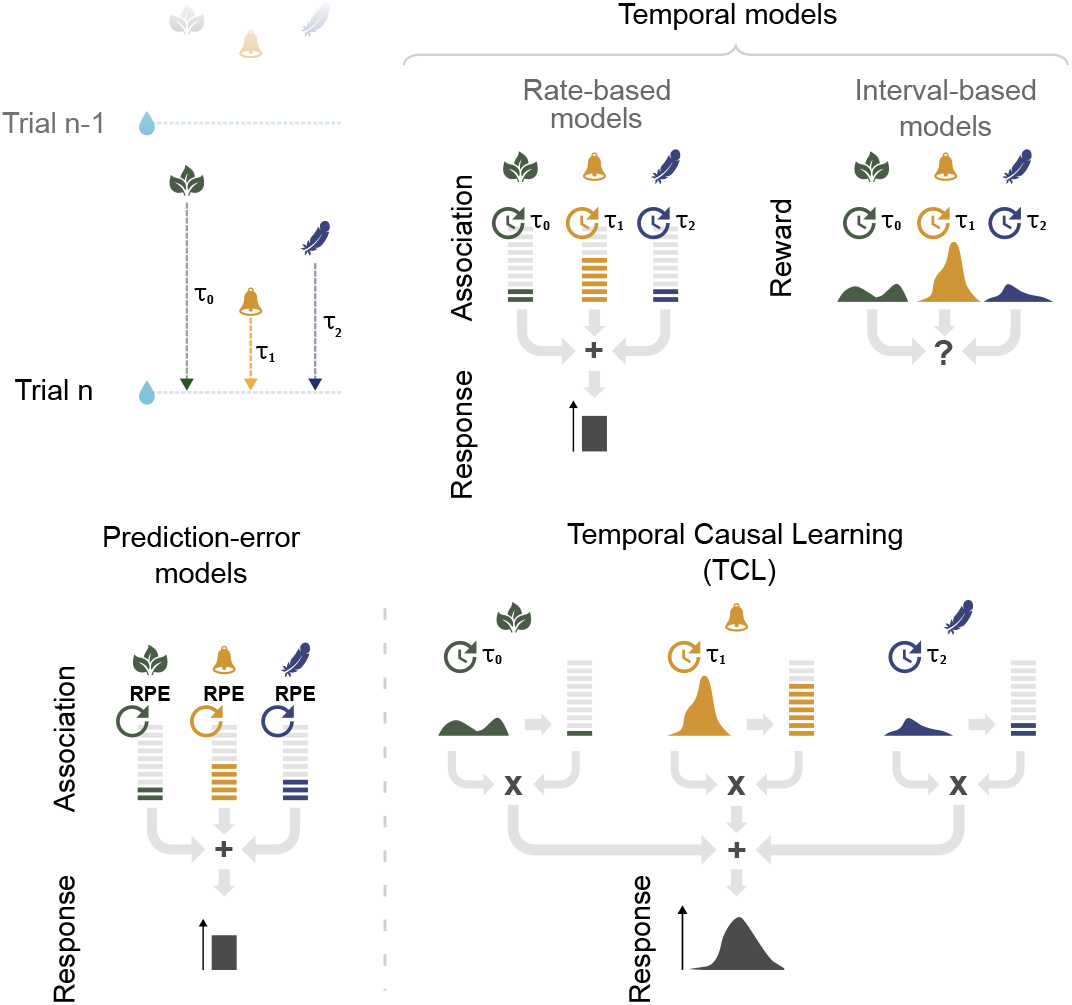
Associative learning theories. We distinguish two large classes of associative learning models: Prediction-error models (bottom left) and temporal models (right). Proposed temporal models can be classified into rate-based models and interval-based models. Rate-based models estimate the rates of cues and rewards to produce a response that depends on the ratios of these quantities. Interval-based models estimate the full distribution between cues and rewards. It is, however, unclear how, in a purely interval-based model, a response is computed. Classical prediction-error models (bottom left) do not estimate time, but update the value of association on each trial based on the difference between the actual reward and the predicted reward. The response reflects the weighted sum of the different associative values. The model proposed in this study (bottom right) is a temporal model that estimates both the intervals between events and a causal association term. The response of the agent reflects the sum of the interval estimates weighted by the causal association. While this model explicitly constructs a representation of time, we show that this model can also be approximated by a prediction-error model.

We predict a neural signal with distinctive learning dynamics that encodes the causal association between the cue and reward. Possible candidates include dopamine itself [21, 48] or other neurotransmitters such as acetylcholine which gates dopamine-dependent learning [49– 51]. In the delay-conditioning protocol, the dynamics of the associative signal are non-monotonic (Figure 6d), with its trajectory over learning influenced by the animal’s prior exposure to rewards. The prediction error term, Δ𝓁_*c*_, associated with this update rule, rises sharply before saturating near zero. The association is acquired during the rising phase. Meanwhile, a parallel set of signals (Δ*w*_*µ*_) updates information about the distribution of cue-reward intervals, and exhibit behavior similar to reward prediction errors (Figure 6a,c) in the commonly used delay-conditioning protocol. Notably, the update associating with timing could be tied to serotonin, which regulates the temporal window during which stimulus-shock associations form in *Drosophila* [52].

Consistent with the dynamics of dopamine observed in [21], the evolution of both RPE signals (Δ𝓁_*c*_ and Δ*w*_*µ*_) in the model is gradual and yet leads to the abrupt emergence of a response. The model predicts that the two prediction error signals display similar dynamics but with opposite signs. One potential approach to disentangle neural correlates of causal learning and temporal coding is to design an experiment in which the cue-reward interval follows an atypical (say bimodal) distribution. Our model predicts that distinct signals will code for different intervals, but a common signal will code for causal associations. Systematically varying the temporal structure (e.g., adjusting the likelihood of one mode in the distribution) and the cue-reward contingency (e.g., changing the proportion of rewards following the cue) would allow for examining whether specific neuromodulators convey timing information or convey causal associations.

The theory does not explain how animals infer causes in realistic scenarios that involve many putative causal stimuli. Attentional mechanisms may play an important role in such scenarios [53, 54]. An attentional mask could be introduced as a scaling factor that modulates the rate at which the stimulus-reward interval is learned based on the probability that the stimulus is causal. Another extension to account for changing environments is to incorporate the influence of context, where each context is associated with a common temporal structure of causes and effects [27, 55, 56]. The Bayesian framework is easily extended to account for higher-order conditioning by introducing the possibility that primary causes of reward may have their own secondary causes. While a full-fledged theory of temporal reinforcement learning incorporating attention, higher-order conditioning, actions and context remains to be fleshed out, this work establishes a link between reward prediction errors and the learning of temporal relationships, and thereby offers a foundational basis for such a theory.

## ACKNOWLEDGMENTS

We wish to thank Randy Gallistel for sharing recent experimental data and for useful comments. GR was partially supported by a joint research agreement between Princeton University and NTT Research Inc. SJG was partially supported by the Air Force Office of Scientific Research grant FA9550-20-1-041.

## IV. METHODS

### A. Data Curation

Experimental data for the plots were used directly from openly accessible datasets whenever available. When datasets were not accessible, data points were extracted from published figures using the tool automeris.io.

### B. Model Simulations

All simulations of the biologically-inspired version of the model (TCL) presented in the main figures of the paper were conducted with the following parameters: learning rate *η* = 1.5 × 10^−2^, *η*^*′*^ = 10^−2^, and *ε* = −20. A notebook containing all the simulations and the code to plot the figures can be found at : https://github.com/noe-hamou/Reconciling_Time_RPE/.

### C. A Bayesian Framework for Estimating Interval Distributions

We present our generative model and discuss algorithms for inference. In this model, there are *C* possible stimuli generated by a point process. Our goal is to predict when the rewarding stimulus, indexed by *r*, will appear next, based on the times at which stimuli have appeared in the agent’s history. We assume that the rewarding stimulus *r* has a single *cause*, which we index as *c*. This causal stimulus *c* can be any one of the *C* stimuli, including *r* itself. The interval between *r* and its cause *c* is drawn probabilistically from a distribution described further below. We denote *ε*_*i*_ as the logarithm of the ratio between the prior probability that stimulus *i* causes *r* and the prior probability that stimulus *r* causes itself. Clearly, *ε*_*r*_ = 0.

We denote the posterior probability that stimulus *i* is the causal stimulus as *π*_*i*_. Following our terminology in the main text, we call *π*_*i*_ the *association* of stimulus *i* to the reward *r*. After observing data, *π*_*i*_ is given according to Bayes’ rule as

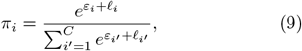

Where 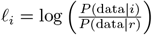 is the log-likelihood of observing the data given that *i* is the cause, relative to the log-likelihood of observing the data given that *r* is the cause. The reward *r* appears after cause *c* with probability *p*, where *p* is drawn from a Beta prior *B*(*p*; *a, b*) with hyperparameters *a* and *b*. If *r* does appear after *c*, the interval *t* is drawn from a distribution *ρ*_*c*_. To sample the interval *t* ∼ *ρ*_*c*_(*t*), we first sample an index *µ* from a Dirichlet-multinomial distribution. Specifically, *µ* (ranging from 1 to *K*) is drawn from a multinomial distribution with class probabilities ***q*** = (*q*_1_, *q*_2_, …, *q*_*K*_). The probabilities ***q*** are in turn drawn from a Dirichlet prior *D*(***α***), where ***α*** = (*α*_1_, *α*_2_, …, *α*_*K*_). Given the sampled index *µ*, we then draw *t* ∼ *ϕ*_*µ*_(*t*), where *ϕ*_*µ*_(*t*) is a normalized probability density function defined below.

We now specify the key features of timescale invariance required to recapitulate the behavioral phenomena of interest. Informally, we would like our model to capture the notion that the animal builds a histogram based on past stimulus-reward intervals. The histogram’s bins are spaced uniformly on a logarithmic scale, and thus the width of a histogram bin is proportional to the bin’s location. Formally,

1. Each class *µ* is associated with a timescale *τ*_*µ*_. The timescales *τ*_*µ*_ are spaced uniformly on a logarithmic scale; that is, *τ*_*µ*+1_ = (1 + *k*)*τ*_*µ*_, where *k* ≪1. The smallest and largest timescales are thus *τ*_1_ and *τ*_*K*_ = (1 + *k*)^*K*−1^*τ*_1_, respectively. We assume that the support of the distribution spans many orders of magnitude, i.e., *τ*_*K*_*/τ*_1_ ≫ 1, which implies *K* ≫ 1 if *k* ≪ 1.
2. The emission probabilities *ϕ*_*µ*_(*t*) for all *µ* have the form 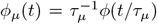, where *ϕ*(*x*) is a density function that normalizes to one, 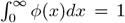. Thus, the functions *ϕ*_*µ*_ tile the interval axis from *τ*_1_ to *τ*_*K*_, and the width of *ϕ* determines how much smoothing is applied when inferring the probability density from finite data. A reasonable choice is to require that *ϕ*_*µ*_ has width proportional to the difference in adjacent timescales, Δ*τ*_*µ*_ = *kτ*_*µ*_. Two possible choices for *ϕ* are: (a) a uniform distribution where *ϕ*_*µ*_(*t*) = 1*/*(*kτ*_*µ*_) when *τ*_*µ*_ ≤ *t* ≤ *τ*_*µ*+1_ and zero otherwise; and (b) a Gamma distribution 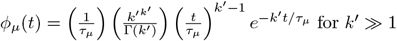 for *k*^*′*^ ≫ 1.

#### Inference

Exact inference in this model is challenging due to the “assignment” problem. For instance, consider a scenario where the causal stimulus *c* (cue) appears thrice at times *t*_1_ *< t*_2_ *< t*_3_, and the target stimulus *r* (reward) appears thrice afterward at times *s*_1_ *< s*_2_ *< s*_3_, with *t*_3_ *< s*_1_. Exact inference would involve iterating over the six possible assignments of the three causal cues with the three rewards. Unless the separation between successive rewards is significantly larger than the cue-reward interval, the number of possible assignments increases exponentially with the number of presentations of *c* and *r* in the worst case scenario. We outline a method for performing approximate inference in Appendix A, although our analysis of the standard conditioning protocol later allows for exact inference due to its trial structure.

### D. Optimal response rates

In this section, we derive the optimal anticipatory response rates (equation (4) in the main text) given the agent’s estimate of the density *p*(*t*) that the reward will appear at time *t*. We set *t* = 0 to be the moment when the most recent stimulus appeared. The agent’s responses are samples from an inhomogeneous Poisson process with a time-dependent, controlled response rate *ω*(*t*). Whenever the agent responds, it incurs a rate-dependent cost *κ*(*ω*). The agent receives reward *R* once it responds after the reward has appeared. We aim to find *ω*^*^(*t*), the optimal response rate that maximizes long-term reward, provided that future rewards and costs are discounted at a rate *λ*. The discount rate motivates the agent to develop an anticipatory response, as the agent would prefer to obtain reward as quickly as possible after it appears.

We first compute the expected discounted reward minus the cost given that the reward appears at time *t*. The expected discounted cost incurred up to *t* is given by 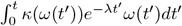. Now, since the reward appears at time *t*, the expected discounted reward is determined by when the agent first responds after *t*. The expected discounted reward is then 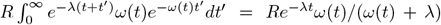. Note that we have ignored the variation in *ω* over the timescale of the first response. The expected discounted cost due to this response is *κ*(*ω*)*e*^−*λt*^*ω*(*t*)*/*(*ω*(*t*) + *λ*). Adding the expected discounted costs and rewards together and averaging over *t*, the net expected discounted reward (including rewards and costs) is given by

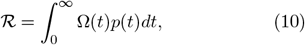

with

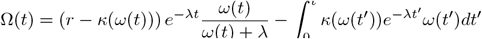

To optimize over *ω*, we take the functional derivative of ℛ w.r.t *ω*(*t*) and set it to zero. This yields an expression for the optimal response rate, *ω*^*^(*t*). Defining 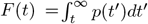, a series of straightforward steps shows that *ω*^*^ satisfies

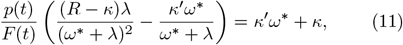

with the constraint *ω*^*^(*t*) ≥ 0. Re-scaling *ω*^*^ with the discount rate *λ*, we get

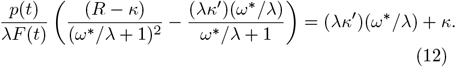

Note that *ω*^*^ always appears as the ratio *ω*^*^*/λ* and the time-dependence only appears as the ratio *p*(*t*)*/λF* (*t*). The optimal response rate thus generically takes the form

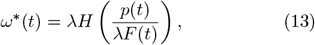

where *H* is a function obtained by solving (12) and depends on the specific form of *κ*(*ω*). If *κ*(*ω*) = *C* (a constant cost), equation (12) gives us

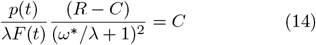

Re-arranging, we get

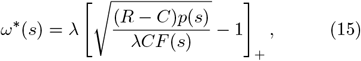

where []_+_ is the rectified linear function. Re-scaling the optimal response rate by *λ*, 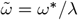, and defining *γ* ≡ (*R* − *C*)*/Cλ* we get

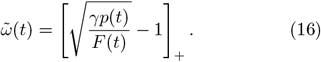

### E. Behavior of the Bayesian model on the delay conditioning protocol

We now examine the behavior of the Bayesian model on the delay conditioning protocol with a single cue and reward. Specifically, the reward is presented *n*_*p*_ times during the pre-conditioning phase with reward-reward interval *C*. Subsequently, the cue and the reward are presented *n* times. The cue-reward interval is *T* and the reward-reward interval remains at *C* (*C > T*). Since *C > T*, the experiment has a trial structure, and *n* indexes the trial number.

Recall that *π*_*c*_ and *π*_*r*_ = 1 − *π*_*c*_ are the associaions of the cue and reward respectively. Our goal is to examine the behavior of *π*_*c*_ as the experiment progresses (increasing *n*) for different experimental protocols (different *n*_*p*_, *C, T*). To derive analytical expressions, we assume *ϕ*_*µ*_(*t*) is a uniform distribution with width *τ*_*µ*+1_ − *τ*_*µ*_ = *kτ*_*µ*_. We do not expect the results to change qualitatively for other scale-invariant choices of *ϕ*_*µ*_(*t*) as long as the density *ϕ*_*µ*_(*t*) is localized around *τ*_*µ*_.

For convenience, we index *π*_*c*_ and 𝓁_*c*_ with the trial number *n* rather than time *t*. From the definition of *π*_*c*_, we have *π*_*c*_(*n*) = *σ*(*ε*_*c*_ + 𝓁_*c*_(*n*)), where *ε*_*c*_ is the relative log prior, 𝓁_*c*_(*n*) is the relative log likelihood after *n* trials and *σ* is the logistic function. The relative log prior *ε*_*c*_ is a constant. We now derive an approximate expression for 𝓁_*c*_(*n*).

To do this, we first derive an exact expression for the likelihood of observing a sequence of intervals *c*_1_, *c*_2_, …, *c*_*n*_. Each of these intervals will fall into one of the *K* bins whose timescales are *τ*_1_, *τ*_2_, *τ*_3_, …, *τ*_*K*_. Recall that the timescales are logarithmically spaced and the width of the *µ*th bin is *τ*_*µ*+1_ − *τ*_*µ*_ = *kτ*_*µ*_. Denote *m*_*µ*_ as the number of intervals amongst *c*_1_, *c*_2_, …, *c*_*n*_ that fall in bin *µ*. Note 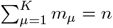.

The likelihood of observing *c*_1_, *c*_2_, …, *c*_*n*_ with a uniform *ϕ*_*µ*_ is proportional to the likelihood of the counts *m*_1_, *m*_2_, …, *m*_*K*_ for a Dirichlet-multinomial distribution. Suppose class *µ* has Dirichlet parameter *α*_*µ*_. Using the expression for the likelihood of a Dirichlet-multinomial distribution, the likelihood of observing *c*_1_, *c*_2_, …, *c*_*n*_ is given by

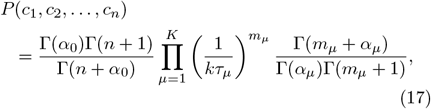

where Γ is the Gamma function and 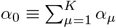.

We now apply the general expression (17) to compute the log likelihoods of observing the cue-reward intervals and the reward-reward intervals in the delay conditioning protocol. We assume *α*_*µ*_ = 1 for all *µ*. This choice corresponds to a uniform prior over the *K* timescales, though the calculation below can be generalized to arbitrary *α*_*µ*_ as long as they are not too large. Since all the cue-reward intervals fall into the same bin (say *µ*_*c*_), we have *m*_*µ*_ = *n* if *µ* = *µ*_*c*_ and 0 otherwise. Moreover, when 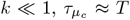. The log likelihood (denoted log *p*_*c*_(*n*)) given *n* cue-reward intervals is thus

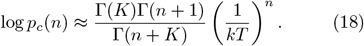

The likelihood of the reward-reward intervals is affected by the *n*_*p*_ prior contextual rewards. Suppose the index of the bin corresponding to the reward-reward interval *C* is *µ*_*r*_. The effect of the prior contextual rewards is to update the Dirichlet prior for the reward-reward interval before the cue-reward pairing begins. Using properties of the Dirichlet distribution, updating the prior is equivalent to updating the Dirichlet parameter corresponding to the *µ*_*r*_th bin from 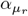 to 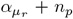 + *n*_*p*_. Assuming again that *α*_*µ*_ = 1 for all *µ*, the log likelihood log *p*_*r*_(*n*) of observing the reward-reward intervals is

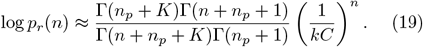

The relative log likelihood 𝓁_*c*_(*n*) = log *p*_*c*_(*n*)*/p*_*r*_(*n*) is then

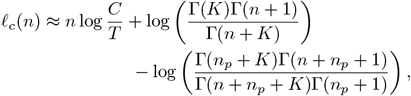

We plot 𝓁_*c*_ for different *n, n*_*p*_, *C/T* values in Figure S1. Next, we find analytical expressions for 𝓁_*c*_ in two relevant limits. Recall that *k* ≪ 1 and *τ*_*K*_*/τ*_1_ = (1+*k*)^*K*−1^ implies *K* ≫ 1 as the minimal timescale *τ*_1_ and the maximal timescale *τ*_*K*_ are separated by orders of magnitude.

Consider *n*_*p*_ = 0. The last two terms in (20) vanish when *n*_*p*_ = 0 and we have 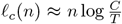. Denote *n*_acq_ the trial at which the likelihood of the data overcomes the prior, 𝓁_*c*_(*n*_acq_) = −*ε*_*c*_. The non-trivial scenario is when *ε*_*c*_ *<* 0, i.e., when the prior probability that the cue is causal is small. When *ε*_*c*_ *>* 0, there is no learning required to form the association. We have

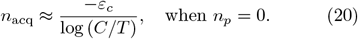

We now consider the asymptotic limit *n*_*p*_, *K* ≫ *n* ≫1. Intuitively, this corresponds to the scenario when the animal has learned the reward-reward interval (to a much better extent than the cue-reward interval) during the pre-conditioning phase and is going through the process of learning the cue-reward interval. Using Stirling’s approximation, we get

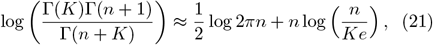

Similarly,

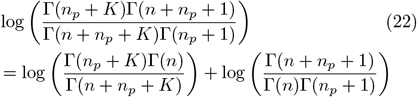

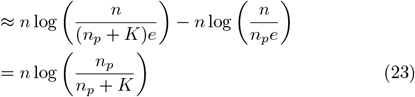

Combining (20), (21), (23), we have

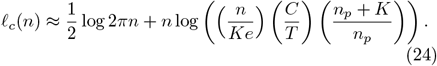

Since the leading order term is *O*(*n* log *n*), we ignore the lowest order term 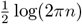 and solve for *n*_acq_. After re-arranging terms, we get

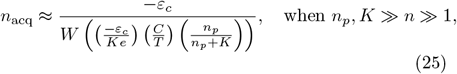

where *W* is the Lambert W function (*x* = *W* (*y*) if *xe*^*x*^ = *y*). From (25), we see that *n*_acq_ has weak dependence on *n*_*p*_ when *n*_*p*_ ≫ *K*. Moreover, *W* (*x*) ≈ *x* for *x* ≪ 1, which leads to

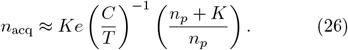

Thus, the 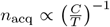 relation indeed holds, but only when the term in the argument of the *W* function in (25) is small.

### F. Other parameterizations of stimulus-reward interval distributions are inconsistent with data

Here, we show that other parameterizations of stimulus-reward interval distributions inadequately explain data. This analysis highlights the importance of our assumption that animals estimate Dirichlet-multinomial distributions (i.e., histograms) of stimulus-reward intervals. We focus our attention on Gamma distributions of intervals:

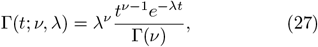

where *ν* is the shape parameter and *λ* is the rate parameter. We assume *ν* is fixed and *λ* is learned. We also consider a scale invariant prior of the rates *p*_0_(*λ*) ∝ 1*/λ* (with the normalization constant determined by upper and lower cutoffs which are not important here). The assumption of fixed *ν* is reasonable as *ν* controls the maximal resolution (i.e. the mean to standard variation) of the distribution after the parameters converge. That is, we assume the animal cannot learn the interval to an arbitrarily high precision. The scale invariant prior *p*_0_ captures timescale invariance but also allows us to derive an exact expression for the data likelihood.

Following our analysis in the previous section, we first write down the likelihood of observing a sequence of intervals *c*_1_, *c*_2_, …, *c*_*n*_ and then specialize to the delay conditioning protocol. The likelihood is given by

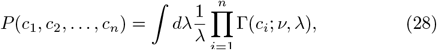

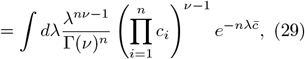

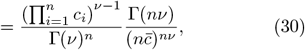

where we have defined 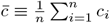 and performed the integral in the second step recognizing that it can be written as a Gamma function.

We use this expression to compute the relative log likelihood 𝓁_*c*_(*n*) as in the previous section. After a few straightforward steps, we get

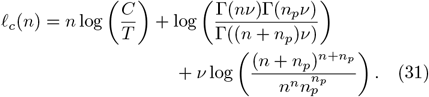

This is an exact expression. We now examine 𝓁_*c*_(*n*) when *n, n*_*p*_ ≫ 1 (recall that in the previous section we assumed *n*_*p*_ ≫ *n* ≫ 1, so this is a weaker condition). Using Stirling’s approximation, we have

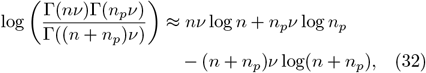

which exactly cancels out with the third term in the exact expression for 𝓁_*c*_(*n*). The leading order term in 𝓁_*c*_(*n*) is thus *n* log(*C/T*). We therefore obtain *n*_acq_ ≈ −*ε*_*c*_*/* log(*C/T*) rather than a power-law scaling observed in data.

### G. The biologically plausible model for timing estimation as an online kernel density estimator

In this section, we show that the biologically plausible model for learning stimulus-reward interval distributions can be interpreted as an online kernel density estimation method. The estimated distribution of stimulus-reward intervals for a particular stimulus is encoded in stimulus-specific weights *w*_*µ*_. Given *w*_*µ*_, the estimated distribution 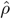 given that the stimulus appears at time *t* = 0 is

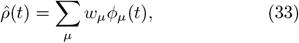

where *ϕ*_*µ*_ is a basis function. We prescribe a update rule for updating weights *w*_*µ*_,

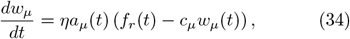

where *η* is a learning rate, *a*_*µ*_ represents the activity of “time cells” (discussed further below) and the constant *c*_*µ*_ is introduced to ensure the weights are normalized. *f*_*r*_(*t*) = ∑_*i*_ *δ*(*t* − *t*_*i*_) is the reward train, where the *t*_*i*_s are the times when the reward appeared. We will show that the kernel estimator is defined by the specific choice of *a*_*µ*_ and *ϕ*_*µ*_. Ensuring that the estimated density integrates to the true probability of reward given the causal cue imposes constraints on *a*_*µ*_ and *ϕ*_*µ*_.

We assume a trial structure. If *η* ≪ 1, integrating (34), the change in *w*_*µ*_ over one trial is

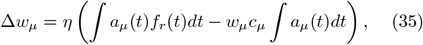

where the integral is over the duration of a single trial and we have used *η* ≪ 1 to ignore changes in *w*_*µ*_ during the trial. To remove the dependence on the second integral, we fix *c*_*µ*_ *=* (∫ *a*_*µ*_ *(t) dt*) ^−1^. This leads to

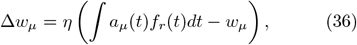

After many trials, the “prediction error” term in the parenthesis converges to zero. After convergence, plugging in *w*_*µ*_ = ∫ *dt*^*′*^*a*_*µ*_(*t*^*′*^) ⟨ *f*_*r*_(*t*^*′*^)⟩ = ∫ *dt*^*′*^*a*_*µ*_(*t*^*′*^)*ρ*(*t*^*′*^) into the expression for 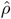 gives

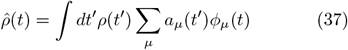

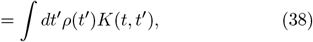

where *K* (*t,t*^*′*^) ≡ ∑_*µ*_ *a*_*µ*_ *(t*^*′*^) *ϕ*_*µ*_ *(t)* is the kernel. The estimated density is thus a smoothed version of the true density.

For any particular choice of *a*_*µ*_ and *ϕ*_*µ*_, we would like to ensure that 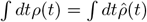, i.e., the true probability of reward appearing within a trial matches the estimated probability. To enforce this constraint, we integrate both sides of (37) to get

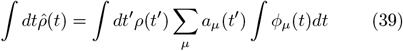

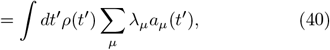

where *λ*_*µ*_ ≡ ∫ *ϕ*_*µ*_(*t*)*dt*. The probability matching constraint (which should apply for arbitrary *ρ*) thus requires ∑_*µ*_ *λ*_*µ*_ *a*_*µ*_ (*t*) = 1 for all *t*. Let’s consider *ϕ*_*µ*_ that are normalized, i.e., *λ*_*µ*_ = 1 for all *µ*, which implies ∑_*µ*_ *a*_*µ*_(*t*) = 1 for all *t*. Recall that in the main text, we specify 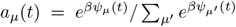 and *ϕ* (*t*) = *ϕ*(*t/τ*)*/τ*, where 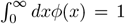. This choice does indeed satisfy *λ*_*µ*_ = 1 for all *µ* and ∑_*µ*_ *a*_*µ*_(*t*) = 1 for all *t*.

Requiring scale invariance would impose additional constraints on *a*_*µ*_ and *ϕ*_*µ*_. In particular, we would like the kernel to be symmetric and translation invariant in logarithmic time coordinates, *x* = log *t*. Suppose the true and estimated densities in *x* coordinates are *ζ*(*x*) and 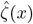, respectively. By the law of density transformations, we have

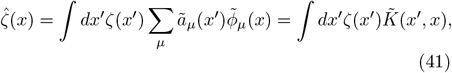

where *ã*_*µ*_(*x*) ≡ *a*_*µ*_(*e*^*x*^), 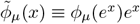 and 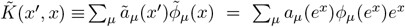. If *ϕ*_*µ*_ is normalized, then so is 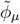. The kernel is symmetric if 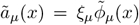 for some constant positive coefficient *ξ*_*µ*_. One can construct appropriate *a*_*µ*_’s and *ϕ*_*µ*_’s by first constructing a symmetric, translation invariant kernel 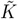 in *x* (on a finite interval) and diagonalizing the kernel to obtain 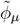’s and *ξ*_*µ*_’s. Our particular choice of *a*_*µ*_ and *ϕ*_*µ*_ (which are derived from eligibility traces *ψ*_*µ*_) is motivated by the simple linear update rule for updating *ψ*_*µ*_.

## Appendix A: Towards an approximate algorithm for Bayesian inference

As remarked in the Methods section, exact inference in the Bayesian model is challenging due to the assignment problem. A dynamic programming algorithm for efficiently computing likelihoods can be derived if we assume that assignments do not “cross.” In the example described in the Methods, this assumption implies that we do not consider scenarios where *c* at *t*_1_ causes *r* at *s*_2_ and *c* at *t*_2_ causes *r* at *s*_1_. We describe the dynamic programming algorithm for *C* = 2, meaning we have two possible stimuli: *c* and *r*.

To perform inference, it is necessary to compute the likelihood of observing the reward *r* at ***s***_*n*_ = (*s*_1_, *s*_2_, …, *s*_*n*_) given that the cue *c* occurs at times ***t***_*n′*_ = (*t*_1_, *t*_2_, …, *t*_*n′*_). We denote this conditional likelihood as 𝓁(*n*^*′*^, *n*) ≡ *P* (***s***_*n*_ | *c* causes *r*, ***t***_*n′*_ ; *p*, ***q***). Once 𝓁(*n*^*′*^, *n*) is computed, the likelihood can be computed from 𝓁(*n*^*′*^, *n*) by averaging over the Beta and Dirichlet priors for *p* and ***q*** respectively.

We compute the likelihood when the latest reward has just appeared so that *t*_*n′*_ *< s*_*n*_. Moreover, since *c* causes *r*, we require that the first *c* appears before the first *r, t*_1_ *< s*_1_. Computing the likelihood of *r* causing *r* is straightforward as a particular *r* is necessarily assigned to its previous occurrence. Note that 𝓁(*n*^*′*^, *n*) = 0 if *n*^*′*^ *< n*: since *r* has a single cause, the number of times *r* appears cannot exceed the number of times *c* has appeared.

Denote Δ_*ij*_ ≡ *s*_*j*_ − *t*_*i*_. We write a recursive expression for 𝓁(*n*^*′*^, *n*):

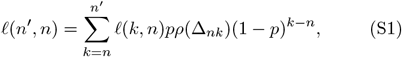

with boundary condition 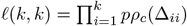, where *ρ*_*c*_ is the cue-reward interval distribution. Note 𝓁(*k, k*) = 0 if any of Δ_*ii*_ ≤ 0 for *i* ≤ *k*. The sum over *k* in (S1) corresponds to the assignment of the latest reward at *s*_*n*_ to one of the cues from *t*_*n*_, *t*_*n*+1_, …, *t*_*n′*_. If *s*_*n*_ is assigned to *t*_*k*_, then by our assumption that assignments cannot cross, all the cues at *t*_*n*_, *t*_*n*+1_, …, *t*_*k*−1_ have not produced a target. The assignment of the *n*th target to *k*th cue leads to the *pρ*(Δ_*nk*_) factor and the fact that the cues from *n* and *k* − 1 have not produced targets leads to the (1 − *p*)^*k*−*n*^ factor.

Equation (S1) provides an efficient update rule for computing the conditional likelihood of observing ***s***_*n*_ given ***t***_*n′*_. However, computing the joint likelihood *P* (***t***_*n′*_, ***s***_*n*_|*c* causes *r*) (after integrating over the priors) also requires computing the likelihood of observing the sequence of *c*, ***t***_*n′*_. The cue sequence ***t***_*n′*_ is either caused by *c* itself or by *r*. Note that we have to assume that the first *c* (if *t*_1_ *< s*_1_) or the first *r* (if *t*_1_ *> s*_1_) has no cause, which otherwise contradicts with the assumption that every cue has a cause that came before it. In general, priors can be integrated out by explicit integration, by substituting the maximum likelihood or maximum a posteriori estimates obtained from gradient descent or Monte Carlo methods.

## Appendix B: Analysis of the TCL model on the delay-conditioning protocol

In this section, we calculate how the number of trials to acquisition *n*_acq_ depends on *C/T* for the TCL model. We will see that the relationship depends on two key parameters: *η*^*′*^, which determines the number of trials averaged over when computing the log likelihood 𝓁_*c*_, and *r*_0_, which is determined by the number of prior rewards *n*_*p*_ presented during the pre-conditioning phase. The update sums the log likelihood over the previous *m* ≈ 1*/η*^*′*^ trials using an exponentially decaying weight. To simplify our analysis, we instead assume a simple unweighted sum over the previous *m* trials.

Since reward-reward intervals and cue-rewards intervals are always *C* and *T* respectively, we need only consider the weights (*w*_*µ*_) corresponding to these two timescales. At the *n*th trial, we denote these weights as *w*_*r*_(*n*) and *w*_*c*_(*n*) for the reward-reward and cue-reward intervals respectively. Whenever reward appears, the weights are updated according to our model as

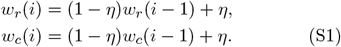

The initial conditions are *w*_*c*_(0) = 0 and *w*_*r*_(0) = *r*_0_, where *r*_0_ *>* 0 when the animal has experienced prior rewards during pre-conditioning.

The log likelihood 𝓁_*c*_(*n*) after the *n*th trial is

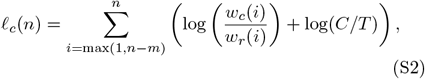

We aim to determine how *n*_acq_ depends on *C/T* for different values of *m* and *r*_0_.

### Case 1

After the weights converge (*w*_*c*_ (∞) = *w*_*r*_(∞) = 1), we have 𝓁_*c*_ (∞) = *m* log(*C/T*). Clearly, learning never occurs if 𝓁_*c*_(∞) *<* −*ε*_*c*_. This condition delineates the regime when the agent never acquires the association. Moreover, if the agent first learns the cue-reward and reward-reward intervals (*w*_*c*_, ≈ *w*_*r*_ 1) and *m* log(*C/T*) *>* −*ε*_*c*_, then from (S2), 𝓁_*c*_(*n*) ≈ *n* log(*C/T*). The agent acquires the association when 𝓁_*c*_(*n*_acq_) ≈ −*ε*_*c*_, and thus, *n*_acq_ ≈ −*ε*_*c*_*/* log(*C/T*).

In the next three cases, we compute *n*_acq_ given that the association is learned before the cue-reward interval is fully learned. Note that the nonlinearity of the association implies that the agent will not show a significant anticipatory response if the cue-reward interval is fully learned (*w*_*c*_ ≈1) but not the association. However, the agent will show a weak but significant response if the association is learned and the cue-reward interval is not fully learned.

#### Case 2

We first consider the case when *r*_0_ = 1 and 1*/η* ≫ *n*_acq_ ≫ *m*. We have

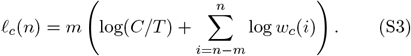

The condition *n*_acq_ ≪ 1*/η* implies that the agent is still learning the cue-reward interval distribution. That is, the weights *w*_*c*_(*i*) are linear in the number of updates: *w*_*c*_(*i*) ≈ *iη*. We get

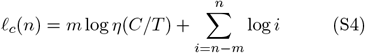

The sum on the right hand side is log *n*!*/*(*n* − *m*)!. Using Stirling’s approximation, this is ≈ *m* log *n*. We then have

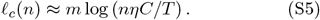

To find *n*_acq_, we solve 𝓁_*c*_(*n*_acq_) = −*ε*_*c*_ to get

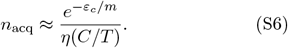

We recover the empirically observed *n*_acq_ ∼ 1*/*(*C/T*) relationship. Let’s denote 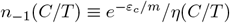.

#### Case 3

We next consider *r*_0_ = 1 but the case when 1*/η* ≫ *m* ≫ *n*_−1_(*C/T*). In this case,

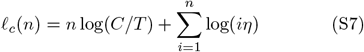

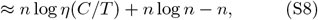

where we have used Stirling’s approximation in the second step. *n*_acq_ is obtained by solving 𝓁_*c*_(*n*_acq_) = −*ε*_*c*_. We have

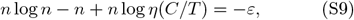

which after re-arranging gives

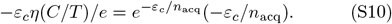

We can express *n*_acq_ using the Lambert W function *W* (*x*),

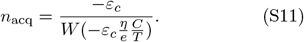

*W* (*x*) ≈ *x* for |*x*| ≪ 1. This implies when −*ε*_*c*_*η*(*C/T*)*/e* ≪ 1, we have

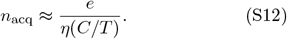

When *W* (*x*) ≫ 1, *W* (*x*) ≈ log *x*, so that when −*εη*(*C/T*)*/e* ≫ 1

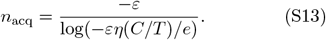

#### Case 4

We now fix *m* such that 1*/η* ≫ *n*_acq_ ≫ *m* and investigate the dependence of *n*_acq_ on *r*_0_. From (S1), we have *w*_*r*_(*i*) = 1 − (1 − *r*_0_)(1− *η*)^*i*^. Since *iη* ≪ 1, we have *w*_*r*_(*i*) ≈ *r*_0_ + *iη*. Plugging this into (S2) for *n*_acq_ ≫ *m*, we get

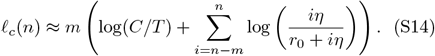

There are now two possibilities. If *r*_0_ ≫ *ηn*_−1_(*C/T*), we then have

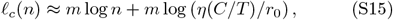

from our previous arguments. This gives

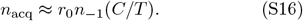

When *r*_0_ ≪ *n*_−1_(*C/T*) (or, say *r*_0_ = 0), the sum in (S14) is negligible and acquisition occurs for *n < m* (recall *m* log(*C/T*) *>* −*ε*_*c*_). Specifically,

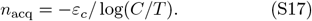

## Appendix C: A brief review of prior models of conditioning

Pavlovian conditioning has a rich history, and various models have been proposed to explain salient aspects of behavioral data. To facilitate clear comparisons, we review these models and their main assumptions. In particular, we will focus on their ability to explain the phenomena associated with timescale invariance. As discussed in the main text, there are two key signatures of timescale invariance in Pavlovian conditioning. First, the response profile after acquisition (measured on probe trails that exclude the US) collapses when re-scaled with the CS-US interval (*T*). Second, the number of trials *n*_acq_ to acquisition has a power-law dependence

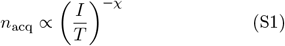

where the exponent *χ* ≈ 1 and *I* is the interval between a US and the next CS. Note that we use the US-US interval *C* = *I* + *T* (also known as the cycle duration) rather than the inter-trial interval *I* throughout the paper. A linear fit (on a log-log scale) to the experimental data compiled in [16] indicates a value of *χ* slightly lower than 1. Equation (S1) suggests that there is no intrinsic timescale when acquisition is quickest; rather the acquisition timescale is set by the inter-trial interval.

### 1. Rescorla and Wagner (1972)

The Rescorla-Wagner (RW) model computes, for each CS (indexed by *i*), the value of the CS-US association *V*_*i*_. The model also considers a “background” cue, which is always present. The RW model also assumes that time is discretized into bins of size Δ*t*. We will use the index *n* to denote the *n*th bin (at time *n*Δ*t*).

The update Δ*V*_*i*_ is proportional to a prediction error signal, which corresponds to the difference between what actually happens, *r* (generally *r* = 1 in the presence of reward, *r* = 0 otherwise), and what is expected (the summed associative value for all current CSs). For simplicity, we consider the case when there is one CS, so that there are two values, *V*_*c*_ and *V*_*b*_, corresponding to the values of the CS and background respectively. The expected reward is

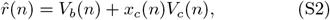

where *x*_*c*_(*n*) = 1 if the cue is present in bin *n* and 0 otherwise. The reward prediction error is 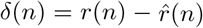 where *r*(*n*) = 1 or 0 depending on whether reward is present or not. The update rule for the value is given by

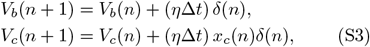

where *η* is the learning rate. The learning rate is often multiplied by a salience factor that depends depend on the CS, which we ignore here.

The response of the agent is a direct reflection of the associative value *V*_*i*_.

The dynamics of the RW model can be obtained analytically in a setting modeled after a delay conditioning protocol. Specifically, suppose that the CS lasts for an interval *T* = *m*Δ*t*. The US appears (as a point event) at the end of the interval. The interval between the US and the subsequent CS is *I* = *m*^*′*^Δ*t*. At the beginning of the protocol, the values are initialized to zero *V*_*b*_(0) = *V*_*c*_(0) = 0. The first CS appears after an interval *I*. From (S3), we will derive an trial-to-trial update rule for *V*_*b*_ and *V*_*c*_. Define 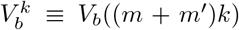 and 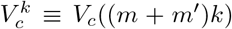. These represent the values after the *k*th US has been presented. A lengthy but straight-forward calculation gives us

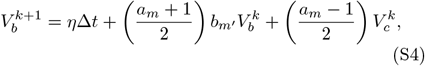

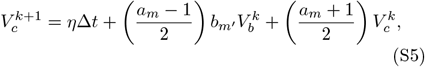

where *a*_*m*_ ≡ (1 − 2*η*Δ*t*)^*m*^ and *b*_*m′*_ ≡ (1 − *η*Δ*t*)^*m*^_*′*_.

Let’s assume max(*m, m*^*′*^)*η*Δ*t* ≪ 1, so that *a*_*m*_ ≈ 1− 2*mη*Δ*t* and *b*_*m′*_ ≈ 1− *m*^*′*^*η*Δ*t*. With this approximation, we obtain

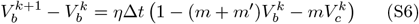

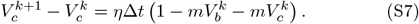

The left hand sides of the above equations are zero at steady state, which gives the steady state values, 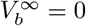 and 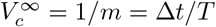. The minimum eigenvalue *λ*_min_ of the matrix

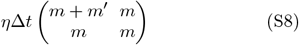

provides an estimate of the number of trials to acquisition. The eigenvalues are

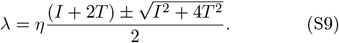

When *I* ≫ *T*, the minimum eigenvalue is *ηT* and the number of trials to acquisition 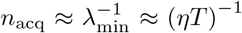 is independent of *I*. The RW model is thus inconsistent with the observed *I/T* scaling. The RW model in this form is applicable for modeling only delay conditioning protocols. Eligibility traces or similar mechanisms have to be included to allow for building associations between temporally separated events. We will consider such models further below.

### 2. Gallistel and Gibbon (2000)

As in our discussion of the RW model, the Gallistel and Gibbon (GG) model [16] considers delay conditioning, where the CS presentation is followed directly by the US, i.e., the CS-US period (*T*) is equal to the CS presentation duration. The period between the US and the subsequent CS is denoted *I*.

The GG model considers timescale invariance of the response profile (known as scalar estimation theory (SET)) and the *I/T* dependence of the number of trials to acquisition (known as rate estimation theory (RET)) separately.

In SET, an animal estimates the CS-US interval 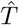 during conditioning. When a CS appears, the animal maintains an “accumulator” or a clock that keeps track of the time since the CS last appeared, say *t*. SET assumes that the response profile is a function of the ratio 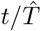. SET further assumes that the estimate 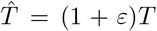, where *εT* is the error in estimating *T*. That is, the error is proportional to the interval itself, similar to Weber’s law. These two assumptions put together are used to explain timescale invariance of the response profiles. SET, however, does not explain *why* we should expect the 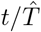 dependence.

RET posits that animals estimate the rate of US with background alone, 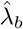, and the cumulative rate due to the CS and background combined, 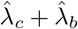. Conditioned response appears when

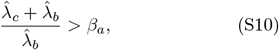

where *β*_*a*_ is a constant decision threshold. The background rate is estimated as 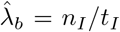 where *n*_*I*_ is the total number of background presentations and *t*_*I*_ is the total exposure to the background. Similarly, the cumulative rate 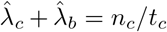, where *n*_*c*_ is the total number of CS presentations and *t*_*c*_ is the total duration the CS was presented. Plugging this in (S10), we get

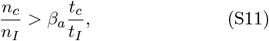

*n*_*I*_ is taken to be one and *t*_*I*_ */t*_*C*_ ≈ *I/T*, which gives for the number of trials to acquisition:

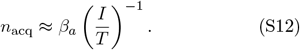

The key assumption (S10) is explained in later papers by invoking an information-theoretic measure of contingency. Specifically, contingency is defined as the difference between the entropy of US arrival with background alone and the entropy of US arrival with background and the CS. It can be shown that for Poisson emission rates, this information-theoretic contingency has the form 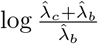 and therefore (S10) is equivalent to the condition that the number of bits that the CS provides about US arrival is greater than a certain threshold.

### 3. Kakade and Dayan (2002)

The Kakade and Dayan (KD) model [23] also considers the delay conditioning protocol considered by the GG and RW models. The KD model was proposed to explain the effect of prior exposure to the US on number of trials to acquisition.

As in the GG model, the model assumes the animal estimates the rate of US during the CS 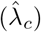 and during the background 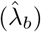. The KD model rather estimates inverse rates (i.e., average interval lengths) using a modified Kalman filter. The filter is set up so as to implicitly satisfy the assumptions of SET. How the intervals are estimated is unspecified. The specific details of the Kalman filter do not affect the rest of the argument and we omit them here. After observing a large number of rewards, the average rates can be simply expressed: 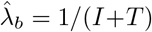 and 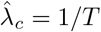. The KD model makes the key assumption that the estimated rates (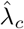 and 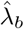) should compete with each other according to their respective reliabilities (*π*_*c*_ and *π*_*b*_ = 1− *π*_*c*_) rather than add as in the GG model. Specifically, in the single cue case, the weighted average 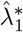 of cue and background rates can be written down as:

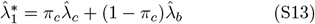

In the case of no cue, the weighted average is:

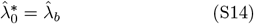

Acquisition is assumed to happen when:

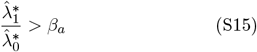

where *β*_*a*_ is a constant threshold. By plugging in the average estimated rates, (S15) becomes:

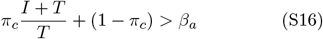

The KD model further assumes that the reliability of the cue has the form

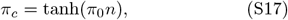

where 1 ≫ *π*_0_ *>* 0. If *n* is small, this can be approximated as *π*_*c*_ ≈ *π*_0_*n*.

Therefore, (S16) becomes:

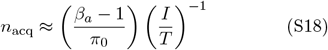

Since we have assumed the estimated rates have converged, there is an implicit assumption here that learning the rates is faster than resolving cue competition.

### 4. Howard et al. (2023)

In this work, Howard et al. [24] provide a model of time-based reinforcement learning that builds upon the experimental observations of the following section, namely that there exists two population of cells, time cells and temporal context cells, that are used to estimate intervals between events. The authors argue that these populations are related by Laplace transforms.

The Howard et al model has similarities with the timing estimation aspect of our model: the temporal context cells show activity that decays exponentially from the previous presentation of a cue (analogous to *ψ*_*µ*_ in our model) and the time cells are most active within a certain interval after the cue (analogous to *ϕ*_*µ*_ in our model). We provide a modified description of the model based on Howard et al, 2023 using notation that is closer to ours, while highlighting key differences.

The temporal context cells encode a Laplace transform of the past timing of a particular cue *c*. We denote 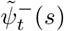 as the activity at time *t* of the temporal context cell associated with this cue and indexed by the inverse timescale *s*. We have

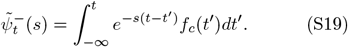

Here *f*_*c*_(*t*^*′*^) is a vector indicating when cue *c* occurred up until time *t* as in our model. The authors assume that *s* is equally spaced on a logarithmic scale, i.e., the spacing Δ*s* ∝ *s*. The 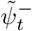’s are obtained using an online update rule similar to our eligibility trace update rule.

Starting with a population of neurons 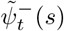, the authors seek to derive the population of neurons encoding a Laplace transform 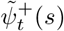 of the future. As we will see below, whenever cue *c* appears 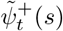 will begin from an initial value and increase exponentially with rate *s*. The initial value depends on a set of weights 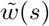 that represent the association between 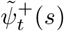 and 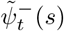. 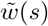 is updated as

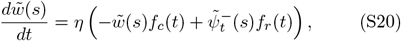

where we have written the above equation in a form similar to our model (see Howard et al 2023 for the original update rule. In particular, we have substituted *ρ* → 1 − *ηdt*).

Let us consider a scenario where the reward follows the cue after time *τ* (a random variable) with probability *p* and does not appear with probability 1 − *p*. At steady state, setting the left hand side to zero and taking expectations on the right hand side, we get

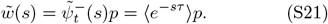

Thus, the association between both events depends on two factors: one measuring whether an event will follow another event, and another related to when the following event will occur.

To update 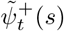, the authors suggest the following update rule (see Howard et al 2023 for the original formulation and argument):

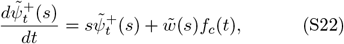

where it is assumed that 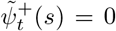 before the cue appears. If the cue appears at time *t* = 0, we have at time *t*,

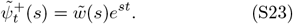

Assuming 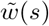 is at steady state, we have

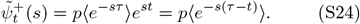

One could recover 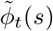 by taking the inverse Laplace transform of 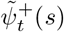. 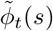 is a direct estimate of the future, and has a form similar to time cells. The authors suggest plausible mechanisms for how the inverse Laplace transform could be implemented in biological networks, including those that use the Post approximation [24] and those that rely on attractor networks [58].

### 5. Cao et al. (2022 and 2024)

In Cao et al. (2022) [45], the authors investigate time cells in the hippocampus, which are neurons that tend to respond after a specific time following an event. These cells correspond to 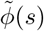 in the Howard et al model from the previous section and *ϕ*_*µ*_ in our model.

In this study, the authors examine how both the duration of their responses (time fields) and the number of active time cells vary with the time since a salient event. Specifically, they find that as the interval lengthens, the time fields extend, and fewer time cells respond. This suggests that these neurons create a compressed representation of time [59]. The authors experimentally test whether this compressed temporal representation by hippocampal cells is consistent with the Weber-Fechner law, which can be stated in this case as:

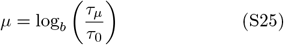

where *µ* is the index of the time cells (sorted by the time of the maximum response), *τ*_0_ is the peak time of the earliest responding time cell, *b* is the base of the logarithm, *τ*_*µ*_ is the time that the *µ*th cell fires maximally. This relationship implies that the difference between two successive peak times can be written as

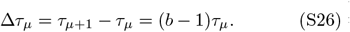

The authors test the hypothesis that the width of the response of individual time cells *σ*_*µ*_ scales with Δ*τ*_*µ*_ to cover all parts of the timeline with the same resolution. Therefore, the first implication of the Weber-Fechner law is:

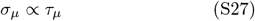

A second implication of the Weber-Fechner law is that the probability to find that the peak time *τ* of a randomly chosen time cell at time *τ* after the cue is:

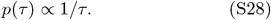

To empirically test the two implications of the Weber-Fechner law, the authors construct a hierarchical Bayesian model of time cell firing. They apply this model to single-unit activity recorded in the dorsal hippocampus of rats during an experiment where the rats were trained to remember the identity of an object over an 8-second interval. The Bayesian hierarchical model estimates the parameters of neural activity at the single-trial, single-cell, and population levels. This approach allows the authors to demonstrate that time field width increases linearly with delay, and that this observation is not the result of a trial-averaging effect. Additionally, they show that the population of time cells is evenly distributed on a logarithmic time scale. They thus confirm experimentally both implications of the Weber-Fechner law.

In Cao et al. (2024) [47], the authors re-analyzed previously published recordings from rodent mPFC neurons [60]. They identified similar “time cells” as those found in the hippocampus, which tile the timeline of future events. Additionally, they identified “temporal context cells” that exhibit decreasing ramping activity from the onset of events, analogous to 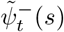 in Howard et al.’s model and *ψ*_*µ*_ in our model. These temporal context cells had been previously described [46]. By fitting both types of cells with a hierarchical Bayesian model, the authors found a continuous logarithmically scaling distribution of exponential rate constants for both past and future cell populations.

### 6. Jeong et al. (2023)

We now consider the model of Jeong et al (2023) [21], called retrospective causal learning theory (RCT). Acquisition in RCT occurs when a CS is identified as a meaningful causal target. A CS is deemed a meaningful causal target when the net contingency (defined below) exceeds a threshold value. RCT applies to trace conditioning, and thus also applies to delay conditioning (the CS onset is considered the stimulus).

The history of an event is maintained as an eligibility trace, which can be expressed at a certain time *t* as

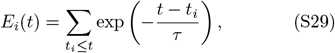

where *t*_*i*_ are the times when the stimulus indexed by *i* is presented and *τ* is the eligibility trace timescale. RCT assumes that an independent (and unspecified) mechanism selects the timescale *τ* = *k*(*I* + *T*) = *kC* proportional to the cycle time. *k* is a positive constant chosen to be 1.2 in [21].

RCT defines the predecessor representation *M*_*ij*_ between an event *i* and a meaningful causal target *j*. The US is *a priori* a meaningful causal target. This representation is updated whenever event *j* occurs using the rule

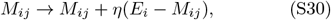

where *E*_*i*_ is the eligibility trace when the US appeared and *η* is a learning rate.

In parallel, a baseline predecessor representation *M*_*i*_ for all stimuli *i* is tracked

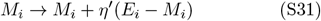

This expression is updated frequently (every Δ*t*) and quickly reaches steady state. The steady state value of *M*_*i*_ keeps track of the average value of the eligibility trace corresponding to event *i*.

RCT then defines the predecessor representation contingency (*C*_*ij*_) linking events *i* and *j*:

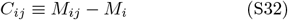

and a corresponding successor representation contingency 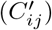

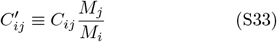

The net contingency 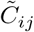 is then defined as a weighted combination of *C*_*ij*_ and 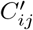:

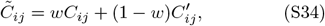

for some weight *w*. RCT posits that a putative causal relationship between *i* and *j* exists if 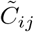 exceeds a threshold *β*_*a*_.

RCT additionally considers an adjusted net contingency, which takes into account the case when *i* itself is caused by another event *k*. We ignore this modification as we will specialize to the delay conditioning protocol (discussed in the previous sections) which only has two events, the CS and the US.

We now examine the behavior of RCT in the delay conditioning protocol considered in the previous sections. We use the subscripts *i* = *c* and *j* = *r* to denote the CS and the US respectively.

In this case, it can be easily shown that *M*_*c*_ = *M*_*r*_ so that 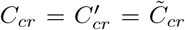. Since *M*_*c*_ is updated frequently, we replace *M*_*c*_ with its average value over a trial:

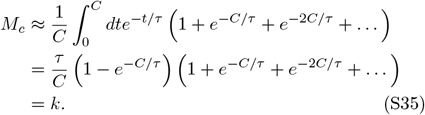

For *M*_*cr*_, after *n* rewards, we have :

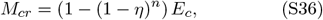

where *E*_*c*_ is the value of the eligibility trace when the US appears, which after summing up over past CS appearances, is given by *E*_*c*_ = *e*^−*T/k*(*I*+*T*)^*/*(1 − *e*^−1*/k*^).

The acquisition criterion is obtained by solving

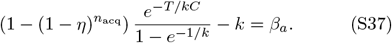

Clearly, the number of trials to acquisition *n*_acq_ depends only on the ratio *C/T*. The *n*_acq_ ∝ (*C/T*)^−1^ scaling however is not reproduced. Moreover, we find that the values of *k* and *β*_*a*_ have to chosen carefully so that acquisition occurs at all.

---

**FIG. S1.**
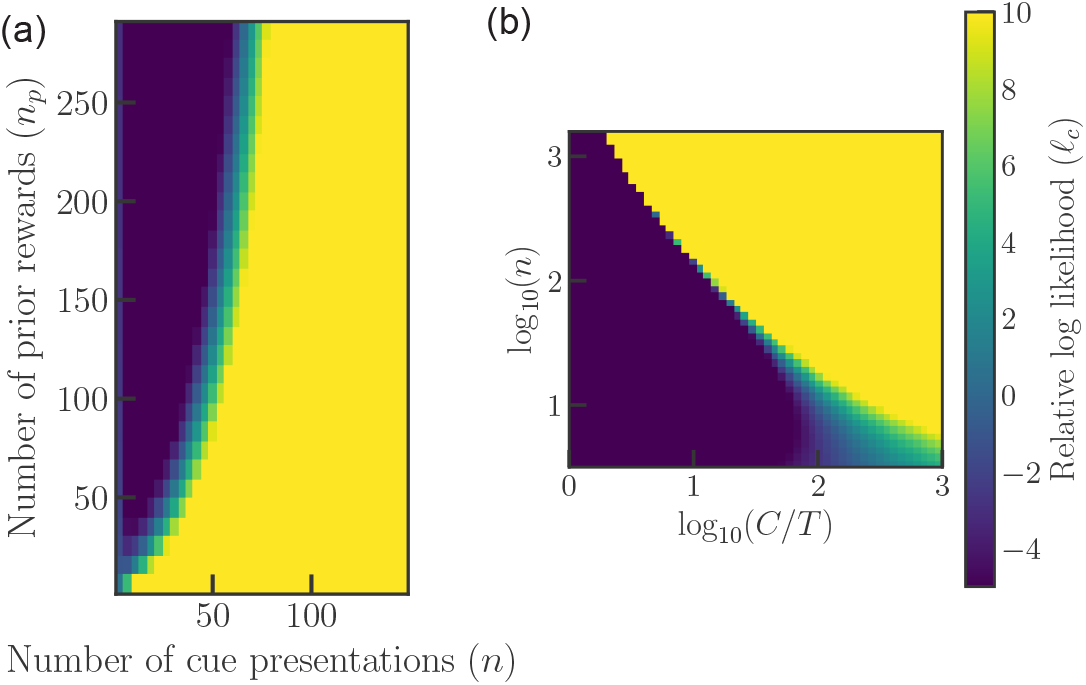
**(a)** Heatmap of the relative log likelihood 𝓁_*c*_ in the Bayesian model for different number of prior rewards (*n*_*p*_) and cue presentations (*n*). **(b)** Heatmap of 𝓁_*c*_ in the Bayesian model for different *n* and *C/T*. Note that *C* is the reward-reward interval and *T* is the cue-reward interval. We fix *C/T* = 4 in panel a, *n*_*p*_ = 5000 in panel b and the number of timescales *K* = 100 in both panels. The colorbar is shared.

**FIG. S2.**
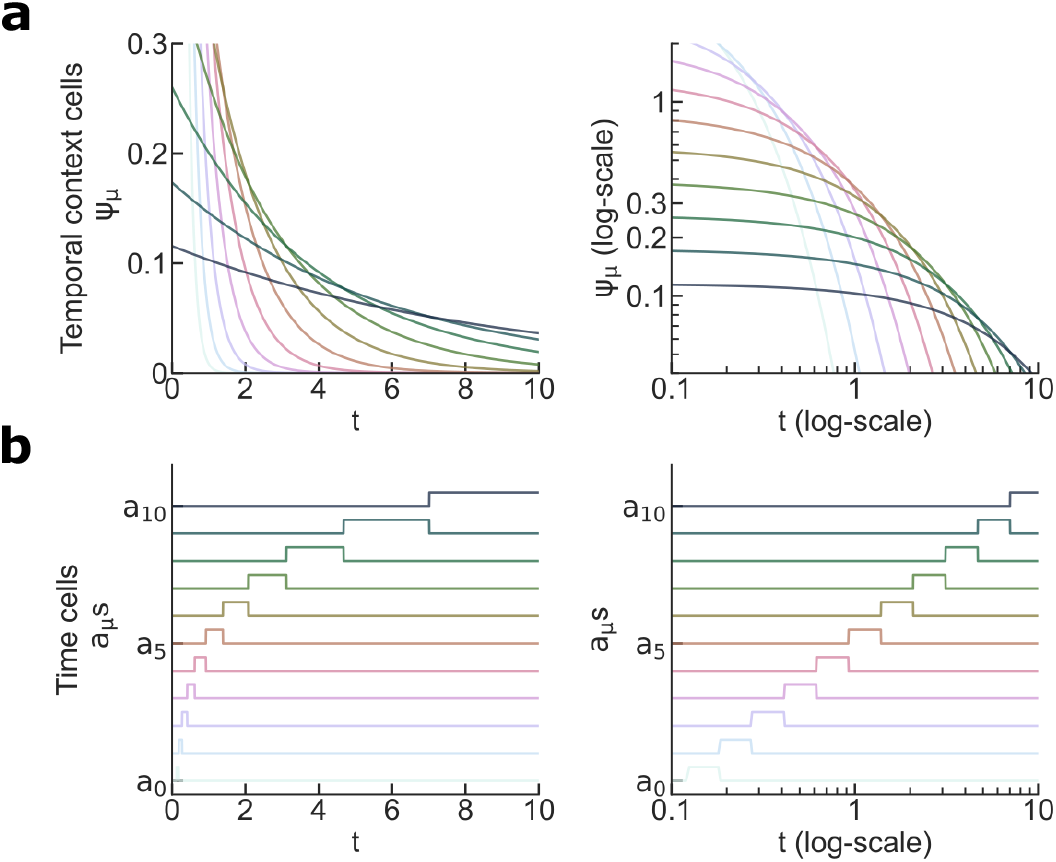
**(a)** Evolution of eligibility traces (*ψ*_*µ*_) for different temporal context cells on a linear time axis (left) and on a logarithmic time axis (right). **(b)** Evolution of the activity of the corresponding time cells (*a*_*µ*_), on a linear (left) and logarithmic (right) time axis. For visualization purposes, time cell activities are spaced vertically.

**FIG. S3.**
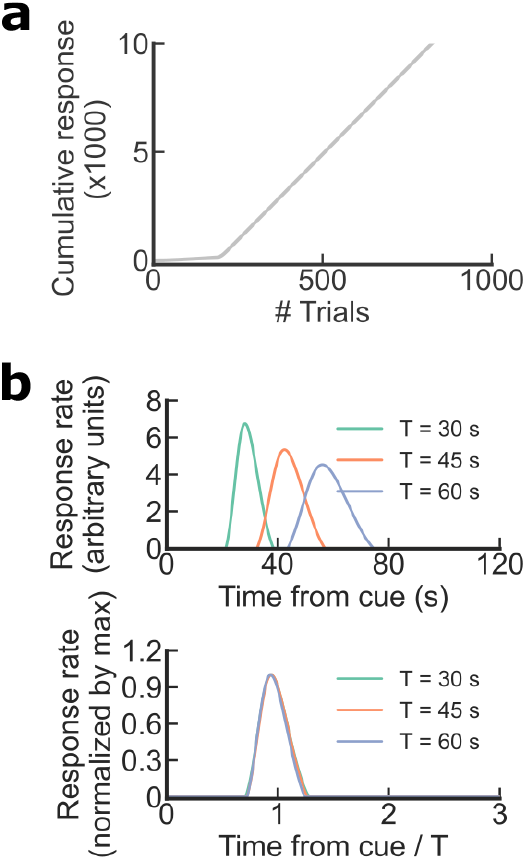
**(a)** Discontinuous Learning curves in the TCL model. Cumulative responses of an agent undergoing a Pavlovian conditioning framework. **(b)** Time-scale invariant responses in the online model. Model responses for different times between cue and reward T (left) and after normalizing the response rate by the maximum response rate and normalizing time by T.

**FIG. S4.**
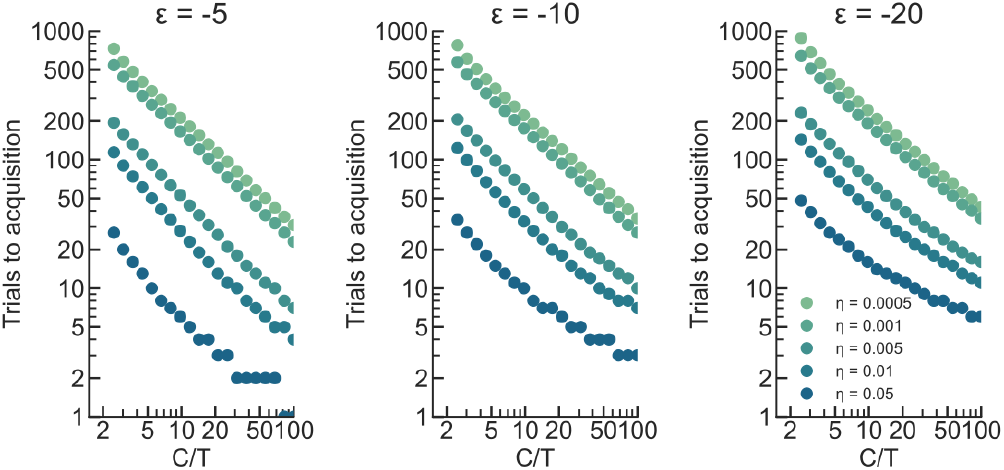
Simulations of the number trials to acquisition for different values of *ε*_*c*_ and learning rates *η*.

